# Tissue flow through pores: a computational study

**DOI:** 10.1101/2021.03.25.436985

**Authors:** Felix Kempf, Andriy Goychuk, Erwin Frey

## Abstract

Cell migration is of major importance for the understanding of phenomena such as morphogenesis, cancer metastasis, or wound healing. In many of these situations cells are under external confinement. In this work we show by means of computer simulations with a Cellular Potts Model (CPM) that the presence of a bottleneck in an otherwise straight channel has a major influence on the internal organisation of an invading cellular monolayer and the motion of individual cells therein. Comparable to a glass or viscoelastic material, the cell sheet is found to exhibit features of both classical solids and classical fluids. The local ordering on average corresponds to a regular hexagonal lattice, while the relative motion of cells is unbounded. Compared to an unconstricted channel, we observe that a bottleneck perturbs the formation of regular hexagonal arrangements in the epithelial sheet and leads to pile-ups and backflow of cells near the entrance to the constriction, which also affects the overall invasion speed. The scale of these various phenomena depends on the dimensions of the different channel parts, as well as the shape of the funnel domain that connects wider to narrower regions.

## 1 Introduction

In their natural environments, collectively moving cell assemblies often face obstacles or barriers. Such external constraints hinder free movement and force the cells to squeeze through narrow gaps or move along externally predetermined routes. Examples for such situations are cancer development and morphogenesis [1, 2, 3, 4, 5], where cells move as cohorts and are surrounded by other cells and extracellular tissue.

To investigate how such extracellular constraints alter cellular behaviour, idealised *in-vitro* experiments have been set up in environments with different geometries [6]. This has been realised for single cells that squeeze through tight capillaries [7, 8], but also in numerous experiments on collective cellular assemblies in diverse settings like straight 2D troughs [9, 10, 11, 12, 13, 14] and 3D tubes [15], winding canals [16], and expanding or narrowing channels [17]. The general question is what phenomenological effects are induced by external confinement and guidance, for example regarding flow behaviour, cell shape, spatial arrangement of cells in the cell sheet, or cell division.

Computational models have been employed to help us gain a better understanding of the essential mechanisms that govern cell behaviour in situations like those described above [18, 19]. The behaviour of cells is governed by numerous complex mechanical and regulatory internal processes as well as chemical and mechanical interaction between cells. Computational models that simulate single cells attempt to reproduce the observed behaviour at small scales, or at least certain aspects of it, by reducing the complex real-world machinery to simpler mechanisms with the aim of capturing the important features of cell behaviour like migration or interaction with other cells. The intention is to construct a minimal functional system that retrospectively justifies the initial guess as to what features were deemed important.

Four types of models for collective cellular dynamics that follow this idea are the so-called Cellular Potts Models (CPM) [20, 21, 22], models in which cells are represented as self-propelled interacting particles [23, 24, 25], phase-field models [26, 27, 28, 29, 30], and so-called vertex models [31, 32, 33, 34, 35]. In the context of cellular assemblies moving in tight channels, self-propelled particle models have been used to investigate straight, converging, and diverging channels [25]. Another approach uses continuum models in which single cells are not resolved. These take into account fundamental symmetries and coarse-grained mesoscopic forces, which provide a larger-scale picture of the system, and they have already been shown to be applicable to cellular systems [36, 37, 38, 39, 18]. Continuum models have also been used to simulate the dynamics of active matter confined to elongated channels [40, 41] and to model invasion of active matter into a narrow straight capillary [42].

In this work, we want to extend previous studies that analysed how confinement in a straight channel affects cellular motion. To that end, we investigate how a cellular sheet adapts to a channel of varying diameter. Similar situations can be expected to arise in real-world scenarios where a migrating cellular cluster is forced to squeeze through a narrower capillary of surrounding tissue.

The paper is organised as follows: In Sec. 2, we explain our setup and the main features of our model for simulating the epithelial layer, focusing on properties that are central to our specific study. Presenting the numerical results in Sec. 3, we first give a qualitative overview and then analyse the cell sheet’s internal spatial order and collective dynamics, as well as the dynamics of individual cells. Finally, we discuss consequences of variations in channel geometry and model parameters before we conclude with a summary and discussion in Sec. 4.

## 2 Setup and model

We investigate the internal spatial organisation and dynamics of an epithelial cell sheet encountering, traversing, and emerging from a bottleneck, whose geometry is shown in Fig. 1A. This setup is intended to be an idealised representation of a capillary-like constriction that cells might encounter in their natural environment. We first let a cellular sheet grow in a reservoir to the left of the channel (not shown in figures) and at *t* = 0 open the channel to initiate invasion. From that point on, we monitor the internal spatial organisation and dynamics of the cellular sheet.

**Figure 1:**
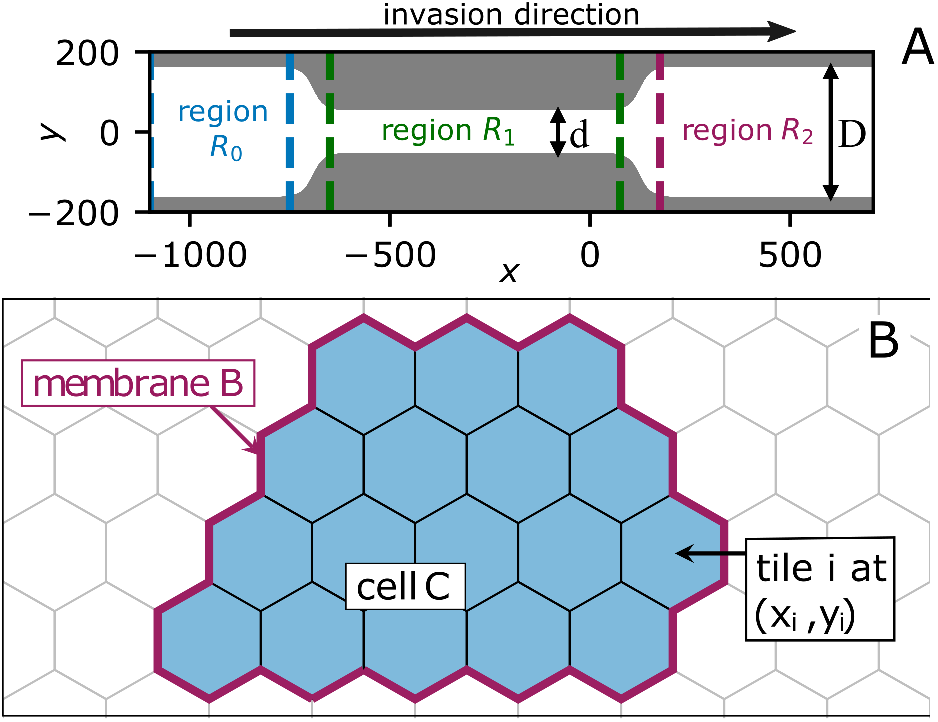
Simulation setup and model. (A) Geometry of simulation setup. Cells enter the channel from a reservoir (not shown) on the left. *D* and *d* denote the diameters of the wider parts and the narrower part (“bottleneck”) of the channel, respectively. Regions *R*_0_, *R*_1_, and *R*_2_ denote the space before, within, and beyond the bottleneck, respectively. In this work, we will analyse these regions separately to investigate the influence of the bottleneck on cell motion. (B) Sketch illustrating the model. On the hexagonal grid, a cell C can occupy several tiles (marked blue). The cell boundary B (violet) is then formed by their outermost edges. Movement of cells is realised by occupying and abandoning tiles, the dynamics of all cells is governed by a kinetic Monte-Carlo algorithm.

In our computational approach, we use a Cellular Potts Model (CPM) that simulates individual, spatially extended cells and their internal dynamics, as well as their mutual interaction on a two-dimensional plane. This computational model adopts an integrative perspective on high-level cell functionality, including the propensity of cells to establish and maintain cytoskeletal polarities, cell-cell and cell-substrate interactions, as well as a basic notion of cell mechanics.

Models of this kind have successfully in reproduced the cellular dynamics of single cells moving in stripe-shaped environments [43] and on substrates of varying stiffness, [44] as well as small numbers of cells in circular [45] or channel-shaped [46] environments, and cell sheets of 2000+ cells expanding into free space [22, 47]. We therefore would argue that the CPM, despite its simple structure, captures essential features of the complex mechanisms that govern real-world epithelial cell dynamics, and permit general predictions of the behaviour of cells under *in-vitro* conditions.

Specifically, we use the model and implementation described in Ref. [22] with slight modifications in the cell-division mechanism (for details see sections A.1–A.3). Figure 1B illustrates how cells are represented in the model. It is a cellular-automaton-type model, where space is discretised into a hexagonal tiling.

A cell occupies several tiles in the plane that reflect the contact area of the cell with the surface and every tile can be occupied by only one cell. The membrane of the cell is formed by the perimeter of the tiles that it occupies. By occupying new tiles or leaving previously occupied ones, cells can change shape and size and, via a combination of several events, migrate on the substrate. The dynamics is governed by a kinetic Monte-Carlo scheme which aims to minimise a global effective free energy. This free energy depends on the collective arrangement of cells, as well as the state of their internal variables, and changes over time as a consequence of the non-equilibrium processes that mimic internal processes of the cells. Among the mechanisms that contribute to the free energy, we will here focus on those that are most relevant for our study:

- A *preferred cell area and membrane length (circumference)*: A term 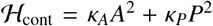 for every cell, which restrains membrane and cortex deformations. Here, *κ_A_* is the area stiffness, *κ_P_* the perimeter stiffness. Together with substrate adhesion, which favours larger cells (as explained below), this leads to a preferred cell size.
- *Cell-cell adhesion* and *cell-cell dissipation*: Two cells occupying neighbouring tiles obtain an energy benefit that reflects the formation of intercellular bonds (cell-cell adhesion), while loss of such mutual contacts and the associated breaking of bonds is energetically penalised (cell-cell dissipation). These effects combined lead to coherent cell structures in the simulation and thus ensure the cohesion of the cell sheet. Dissipation also leads to “viscosity-like” effects.
- *Interaction with the underlying substrate regulated by an internal polarisation field*: The cytoskeleton of the cell, which is regulated by networks of signalling proteins [48, 49], adheres to the substrate via integrins and determines the tendency of a cell to form protrusions or to retract. The contribution to the Hamiltonian that is associated with cell shape, 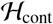 is purely contractile, and thus would, by itself, lead to detachment of cells from the substrate thereby favouring cells with zero contact area. To counteract this cell contractility and detachment, we use an effective surface adhesion energy, the scalar polarisation field ϵ. This polarisation field serves as a proxy for the ability of cells to form focal adhesions and to facilitate polymerisation of the actin cytoskeleton. A higher value of the polarisation field can be interpreted as a stronger local adhesion, therefore counteracting retractions and facilitating protrusions. Similarly, a low value of the polarisation field means that it cannot locally balance out cell contractility, so that the cell will retract (i.e. detach from the substrate). In turn, we represent the signalling networks that regulate the formation of focal adhesions and polymerisation of the cytoskeleton via two prototypic feedback loops: in the vicinity of protrusions (retractions) we increase (decrease) the polarisation field. It is this feedback loop that prevents the system from relaxing to a time-independent global minimum-energy state and induces an internal polarisation and persistent motion of cells. The interplay of the polarisation field and cell-cell adhesion results in collectively migrating, coherent patches of cells.
- *Internal cell-cycle*: To achieve invasion of cells into the capillary, proliferation of cells needs to be implemented. After a growth period in which *κ_A_* and *κ_P_* are continuously adapted such that the preferred size of the cell doubles over the full period, the cell enters a division phase. The polarisation field uniformly drops to zero and at the end the cell splits into two identical daughter cells. After division, the daughter cells are in a quiescent state, i.e. there is no growth. After a certain time, cells are ready to enter the growth phase again, but do so only if the cell size, via fluctuations, exceeds a certain threshold. The latter effect ensures that growth is halted above a certain cell density. In our simulations, cell division is limited to the reservoir, as we assume that the entire trajectory in our system occurs on timescales in which cell division can be neglected.

For a complete listing of the parameters and their values used, see section A.

## 3 Results

### 3.1 Velocity and density profiles

The bottleneck with its smaller diameter *d* hinders free homogeneous expansion of the cell sheet in the direction of the channel axis. As illustrated in the example in Fig. 2, the constriction affects cell velocities as well as the number density. In this example, *D* = 3*d*, where *d* corresponds to roughly the width of 10 cells. The length of the bottleneck is ≈ 5*d*.

**Figure 2:**
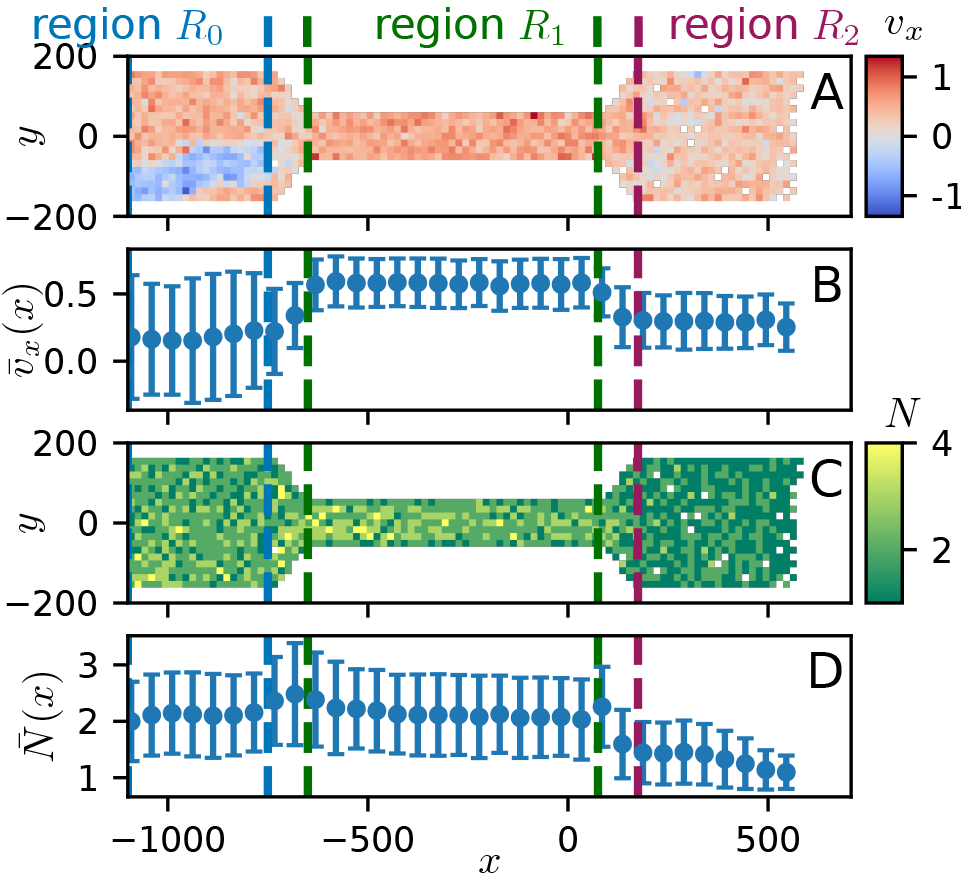
Example of a simulation in a constricted channel. Bottleneck diameter d corresponds to about 10 cells, diameter of the broad regions is D = 3d, length of bottleneck ≈ 5d. (A) Heatmap showing cell velocities along the channel axis *v_x_*(*x, y*) at a late time-point in the simulation. (B) *v_x_*(*x, y*) averaged over *y* and time. At the average velocity of ~ 0.5 in the narrow region *R*_1_, a cell takes ~ 30 time steps to cover the distance corresponding to one cell diameter. (C) Heatmap showing the density distribution *N*(*x, y*) at a late time-point in the simulation. (D) *N*(*x, y*) averaged over *y* and time. Observables in B and D show higher values in region *R*_1_. Error bars in B and D show the standard deviation.

The forward velocity *v_x_*(*x, y*) (Fig. 2A,B) increases in the narrow region and drops again following passage through the constriction to a value larger than the velocity at the entrance. This disparity of velocities is explained by clusters of cells that are forced to retreat owing to the congestion caused by the constriction - a phenomenon we call “backflow” - which leads to a reduction in the average 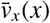 and an increase in the standard deviation (Fig. 2A). This is in contrast to what would be observed for laminar flow of an incompressible fluid where the fact that the channel width is the same on both sides of the narrow region would lead to identical values of 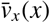 [50].

In comparison, the number density *N*(*x, y*) of cells (Fig. 2C,D) is roughly constant in the regions *R*_0_ (ahead of the bottleneck) and *R*_1_ (inside the bottleneck), but shows a slight rise right in front of the funnel and drops considerably upon leaving the bottleneck, i.e. when the channel widens again. The rise points to an accumulation and corresponding compression of cells in the funnel. Interestingly, the higher density of cells is not “translated” into the narrow channel section, but causes cells to change direction, thus causing the backflow. In contrast, the widening at the end of the channel allows expansion of the epithelial sheet, leading to a lower density, or larger cell areas, and reduced velocity in the region beyond the bottleneck. Thus, both findings show that cells do not pass the narrow part smoothly, like a laminar fluid would do, but that the rightwards motion of the cells is distorted and the flow behaviour of the cell sheet clearly shows properties of a compressible fluid. In general, it should also be noted that fluctuations in both observables are quite high, as the standard deviation is of the same order of magnitude as the mean values (Fig. 2B,D).

### 3.2 Spatial organisation

To further assess the effect of the bottleneck (Fig. 1A) on the spatial organisation of cellular tissue, we measured the radial distribution function (RDF) in our simulations. The RDF is defined as

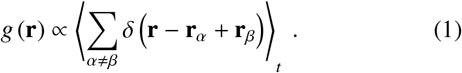

The double sum in Eq. (1) runs over all pairs of cells in the area under observation, with **r**_*α*_, **r**_*β*_ being the positions of the cells [51] while the angular brackets indicate a time average which is taken late in the simulation such that there are cells in all channel regions. In essence, this yields a histogram of the distribution of mutual particle-particle distances **r** that characterises the close-range order of the cells. To analyse the differences between the three regions of the bottleneck setup, we calculate the RDF separately for each.

Figure 3 shows a comparison between a system with a uniform channel width *D* (X1) and a channel geometry which also contains a bottleneck of diameter *d* (X2). In the absence of a bottleneck, the RDF exhibits a set of six fairly pronounced peaks regularly spaced on a circle around the centre. Note that even higher order peaks are clearly visible, indicating a finite range of lattice-like order with hexagonal symmetry. This is reminiscent of a similar degree of ordering observed in two-dimensional colloidal systems [52]. The appearance of such sixfold coordination in cellular sheets is consistent with previous studies, as it was observed very early *in vivo* [53], as well as more recently in Voronoi [54] and vertex [55] models. It can also be motivated from purely topological considerations [56].

**Figure 3:**
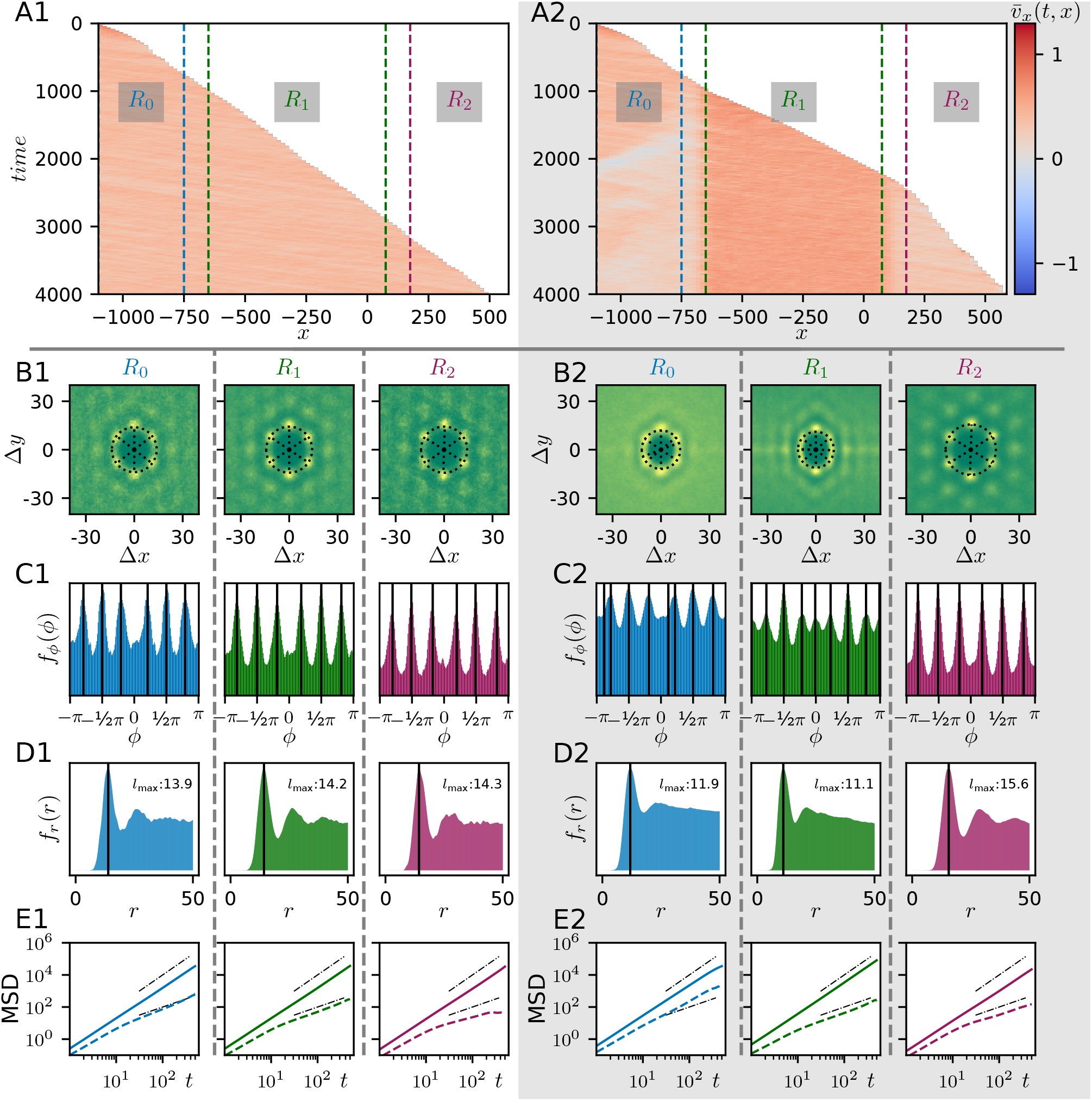
Comparison of cellular order and dynamics between a homogeneous and a constricted channel, channel parameters as in Fig. 2. Left (panels X1): homogeneous channel, right (panels X2): channel with bottleneck. (A) Kymograph of the velocity in channel direction, *v_x_*(*t, x*). Dashed lines indicate transition regions in A2, and are indicated for comparison in A1. Two features of cell motion in the bottleneck configuration (on the right) are clearly visible: the higher velocity in region *R*_1_ (for definition of regions see panel A and Fig. 1) and the backwards motion of cells in front of the bottleneck. Both are already discernible in Fig. 2A,B. (B-D) Radial distribution functions (RDF) for the different regions of the channel, as defined in the main text. The six regularly spaced peaks around the centre for the straight channel in B1 reflect the disposition of cells on a hexagonal lattice, while the more washed-out patterns in regions *R*_0_ and *R*_1_ in B2 show that spatial constraints prevent the cells from adopting that arrangement. For more detailed analysis, the RDF is integrated over the radius (panels C) or the polar angle (panels D). Vertical black lines in C and dashed radial lines in B mark maxima in the marginalised angular distribution function *f_ϕ_*(*ϕ*) as detected by our algorithm (for details see section B.2). Largely unperturbed hexagonal order is reflected by six pronounced maxima, while additional peaks appear at −*π/π* and 0 in region *R*_1_. The vertical black lines in D and the dashed circle in B mark the respective maxima of the marginalised radial distribution functions *f_r_*(*r*) which give the typical nearest-neighbour distances. (E) Mean-squared displacement (MSD, solid line) of cells and cage-relative mean-squared displacement (CR-MSD, dashed line). Black dashed-dotted lines show linear and quadratic scaling, respectively. In contrast to the ballistic MSD, the CR-MSD clearly scales with a lower, but still finite exponent. This shows that cells, unlike particles in a true solid, are not caged, but behave more like diffusive particles suspended in a liquid. Such a combination of properties that is typical for glasses and viscoelastic materials. As an exception, the CR-MSD is superlinear in region *R*_0_ of the bottleneck channel. Here the backflow leads to enhanced relative motion of cells.

In contrast to the fairly regular and homogeneous patterns that we observed in the straight channel, the degree of ordering shows significant spatial variation across the different regions in the presence of a narrower central section. In front of the bottleneck (left panel in B2), the six peaks are blurred in the angular direction and there are no further peaks beyond the closest six. Both observations hint at a distortion of the regular hexagonal order. Inside the bottleneck (centre panel in B2), two additional peaks appear at Δ*y* = 0 which come from neighbours before and after the cell along the channel axis. We suppose that this longer-ranged ordering is due to the fact that the closer channel boundaries force cells towards the centre, and thus hinder the free development of the hexagonal pattern in lateral direction. To the right of the bottleneck, the ordering is fairly similar to that observed for systems with a uniform channel width. This is because, as in the straight channel, cells can invade the free space beyond the bottleneck without hindrance.

In order to bring out the characteristic features of the RDF more clearly, we look at the marginalised angular distribution *f_ϕ_*(*ϕ*) and the marginalised radial distribution *f_r_*(*r*). These represent the integral over the radius and over the polar angle, respectively (for details see section B.2). *f_ϕ_*(*ϕ*) allows one to compare number and arrangement of the maxima in the RDF, *f_r_*(*r*) can be used to determine their typical distance from the centre. The results are shown in Fig. 3C,D.

In the cases where the radial distribution function *g* (**r**) shows six distinct peaks, there are also distinct peaks in *f_ϕ_*(*ϕ*). In contrast, when the RDF is more blurred, then the peaks in *f_ϕ_*(*ϕ*) are also less pronounced. This is especially prominent in the regions *R*_0_ and *R*_1_ for the channel with a bottleneck. In all cases, however, the radial distribution *f_r_*(*r*) shows a distinct peak at position *l*_max_ indicating the typical nearest-neighbour distance. Additional maxima are due to second and further nearest-neighbours. While this inter-cell distance is more or less constant along a straight channel, under the influence of the bottleneck geometry it is considerably larger in region *R*_2_ beyond the bottleneck compared to the other regions *R*_0_ and *R*_1_. These observations from the marginalised distributions *f_ϕ_*(*ϕ*) and *f_r_*(*r*) support the following conclusions: cells are compressed ahead of the funnel and in the narrow region, which translates into a reduced neighbour-neighbour distance, but also induces asymmetries in the hexagonal arrangement.

### 3.3 Collective cell dynamics

In order to assess the dynamics of the cells, we monitored both their collective dynamics (Fig. 3A) as well as the dynamics of individual cells (Fig. 3E).

For the collective dynamics, we follow the time evolution of the mean velocity-profile in channel direction 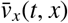 as it is shown in the kymographs in Fig. 3A. Here, the velocity profile for one instant in time is plotted along the horizontal axis using a colour code, and time evolution thus proceeds along the vertical axis from top to bottom. Locations where there are no cells appear white, so that the trajectory of the advancing front of the cell sheet appears as the line that separates the white from the red area. Figure 3A shows that the velocity field is more homogeneous in the straight channel, as there is no backflow or regional velocity increase, and only slight wave-like modulations appear. These modulations are also present in the case with a bottleneck, but in general appear only as slight variations compared to the other effects, such as backflow and velocity increase in the narrow region. Also, we note that backflow appears with some delay after the first cells have passed through the funnel and begun to invade the narrow region.

To characterise the dynamics beyond the mean velocity profile, we next have a look at the second moment of the cell trajectories. A typical measure for the second moment is the mean squared-displacement (MSD), which tells us about the movements of the cells relative to the laboratory frame, i.e. the surrounding confinement, but does not inform us how particles move relative to each other. To quantify this relative motion, we also consider the so-called cage-relative MSD (CR-MSD). In this quantity, the mean trajectories of all neighbours of a cell are subtracted from the cell’s trajectory. Then, the MSD is calculated from this relative trajectory [57, 58] (details in section B.2). In two-dimensional colloidal systems, the CR-MSD, due to its local nature and hence independence from large-scale (Mermin-Wagner) fluctuations, can be used to discern solid and fluid states with more contrast than the ordinary MSD. In a solid, single particles are “caged”, i.e. trapped between their neighbours and thus the CR-MSD is bounded. In contrast, in a fluid, the CR-MSD would be unbounded and would steadily increase with time [58].

In our system, the “ordinary” MSD (solid lines in Fig. 3E) scales with *t*^2^, reflecting the polarised and coordinated motion of the particles through the channel. The CR-MSD (dashed lines in Fig. 3E) in comparison shows a reduced scaling as overall rightwards motion is subtracted, but is not bounded. This indicates that cells are not trapped between neighbours, but that instead there is relative motion between cells inside the sheet and change of neighbours. This dynamic feature, combined with regular hexagonal order, is typical for glass-like systems [58]. Such parallels between glasses and cellular systems have previously been reported in various experimental and theoretical studies. [59, 60, 61, 62, 63]

### 3.4 Varying channel widths

After learning how the presence of a bottleneck affects spatial organisation and spatio-temporal dynamics of cell sheets, we next ask how the observed effects depend on the exact channel geometry. We start by varying the values for the diameters *D* and *d* for broad and narrow channel regions, respectively.

In Fig. 4, we compare the spatial organisation of the cellular aggregate under variation of 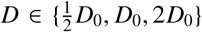 and relative to that 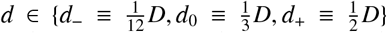, where (*D*_0_, *d*_0_) refers to the parameter combination studied in the previous sections. Comparing the results across all combinations of these parameters, the following general observations can be made. In region *R*_2_ (behind the bottleneck), there is pronounced hexagonal order in all cases. In contrast, in the other regions the degree of spatial ordering depends on the absolute and relative magnitude of the width of the broader sections *D* and the bottleneck *d*.

**Figure 4:**
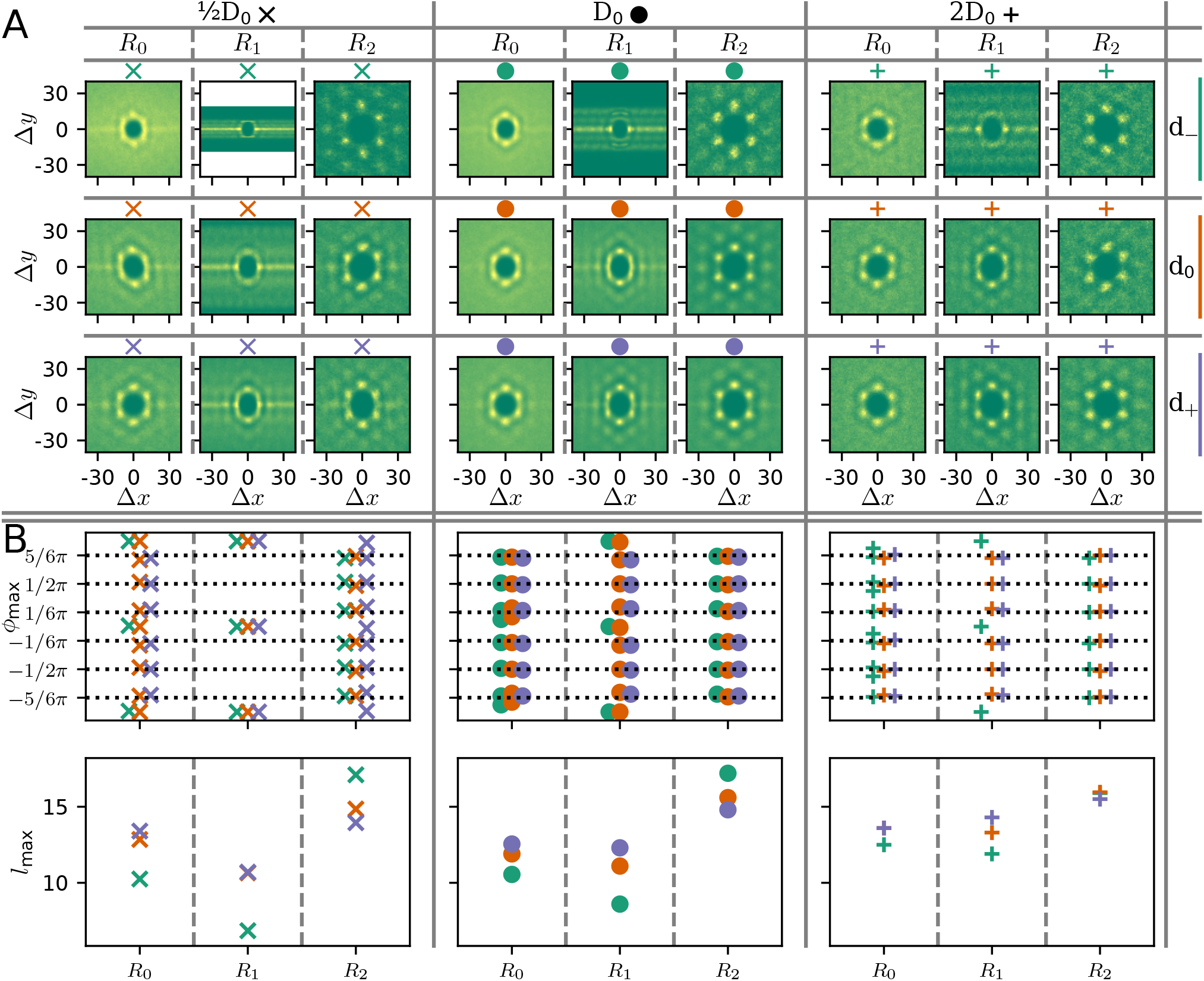
Comparison of cellular order in constricted channels of varying width. (A) Radial distribution functions (RDF) for different diameters *D* of broader channel sections and *d* of constricted channel section (for definition of regions see Fig. 2). From left to right: increase in 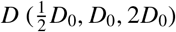, from top to bottom: increase in *d* relative to 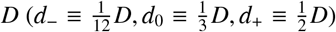. The larger the diameter of the region under consideration, the more the RDF resembles a regular hexagonal configuration and the less order in region *R*_0_, upstream of the constriction, is influenced by the bottleneck. For white areas there is no data due to the very narrow channel. (B) Results of quantitative evaluation of the results in panel (A). The positions of the markers along the horizontal axis of the respective plots correspond to the regions indicated. Varying marker forms indicate varying *D*, varying colors indicate variations in *d* as the small symbols above the plots in part A show. In both panels, plots show results for increasing *D* from left to right. *Upper panel*: Positions of maxima in the marginalised angular distribution function obtained from radial integration of the RDF (compare part C of Fig. 3, and Supp. Fig. 3). For situations similar to those in region *R*_1_ for (*D*_0_, *d*_0_) or 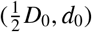, the detectability of the additional maxima depends on the parameters chosen for the maxima detection algorithm (section B.2)), but still reflects the influence of the greater proximity of the walls. *Lower panel*: Positions of the maxima in the marginalised radial distribution function obtained from the angular integral of the RDF. As shown in panel D of Fig. 3, angular integration of the RDF leads to a distribution whose first peak shows the typical nearest-neighbour distance. Variation across regions is the same as that across parameter combinations except for largest (*D, d*). (leftmost plot: violet cross on top of the orange cross for region *R*_1_, same for the + in *R*_0_ in the rightmost plot. Also there in *R*_2_, the orange + is on top of the green one.)

On the whole, there is a tendency that the larger the diameters *D* or *d*, the lower the perturbations of the spatial order, i.e. the closer the ordering is to a regular hexagonal configuration. We attribute this to the fact that for smaller *D* and bottleneck diameter *d*, the bottleneck poses a greater hindrance to free movement of cells. For example, the tight bottleneck *d*_−_ corresponds to the width of 2-3 cells for *D* = *D*_0_. Thus, for extreme constrictions *d*_−_, cells are forced into a single-file-like configuration, as the angular locations of the RDF’s peaks in the upper panel of Fig. 4B indicate. Analogously, the lower panel of Fig. 4B shows that for smaller *d*, the typical distance between the cells, *l*_max_, drops in region *R*_1_ indicating a higher cell density. For region *R*_0_ maxima in the RDF are smeared out in Fig. 4A. That indicates distorted hexagonal order in this region (see also Supp. Fig. 3). These observations show that a bottleneck diameter *d* that permits passage only to a few cells in parallel qualitatively changes spatial organisation inside the bottleneck.

We also analysed how the dynamics of cells is affected by changes in the geometry of the confinement; see the MSD and CR-MSD shown in Supp. Fig. 4 and the corresponding scaling-analysis in Fig. 5. Beyond the bottleneck (region *R*_2_), where as we have seen cellular organisation is hexagonal and cell density is minimal, the MSD indicates ballistic cell movement, while the CR-MSDs’ scaling exponent is significantly reduced and hints at diffusive motion of cells relative to their neighbours. This is the same result as obtained for a straight channel without a funnel (see Supp. Figs. 5–9). In contrast, in the spatial domain ahead of the bottleneck (region *R*_0_), backflow and congestion effects lead to a distinct scaling of the MSDs: especially for smaller diameter *D* of the broader section, ballistic motion is inhibited. At the same time, relative motion as measured by the CR-MSD is superdiffusive and scales comparably to the ordinary MSD. Together, these derivations reflect less directed movement, but enhanced mixing of the cells.

**Figure 5:**
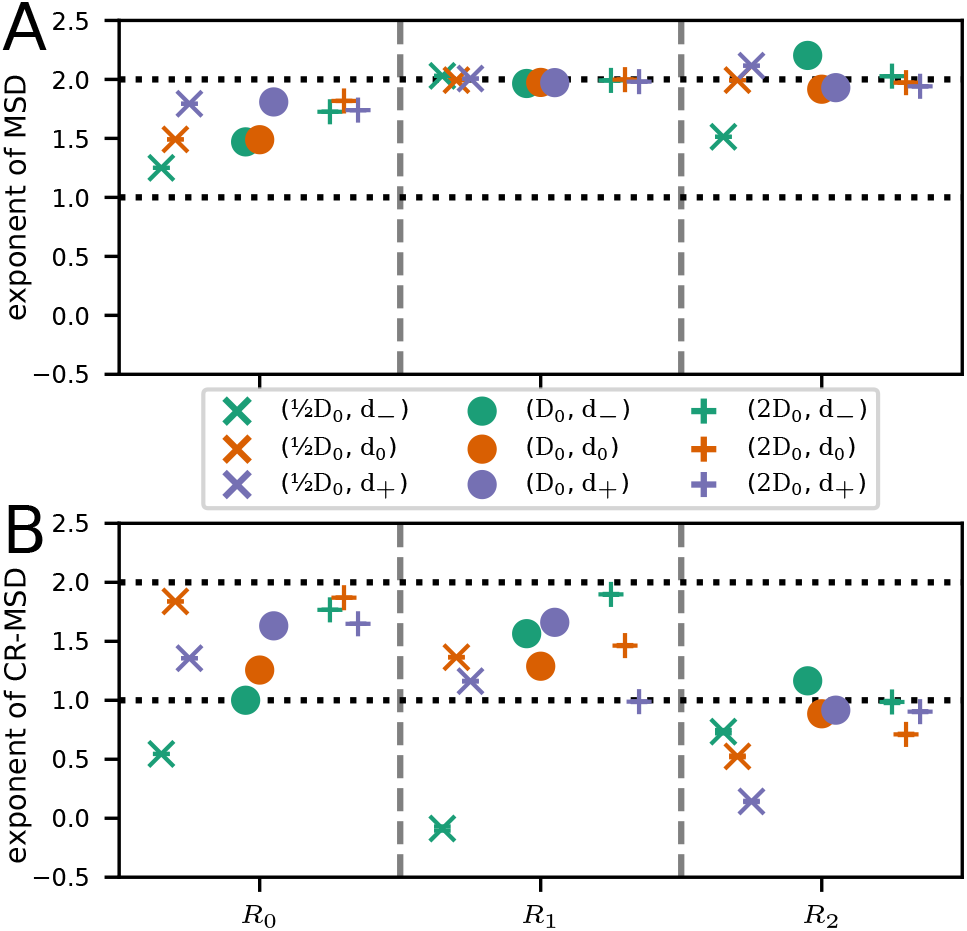
Comparison of MSDs in constricted channels of varying width. (A) Exponent from monomial fit to the MSD for same parameter configurations as in Fig. 4, depicted separately for different bottleneck diameters *d* and regions *R*_0_, *R*_1_, and *R*_2_. Except for the very narrow configuration (blue diagonal crosses), the MSD reflects the ballistic motion of the polarised cells in regions *R*_1_ and *R*_2_. In region *R*_0_, congestion at the bottleneck entrance leads to a reduced exponent. (B) Corresponding exponents from monomial fits to the late CR-MSD (for definitions see section B.2). Compared to the ordinary MSD in A, the exponents are reduced in regions *R*_1_ and *R*_2_, indicating reduced relative motion and thus coordinated motion of cells.

Finally, inside the bottleneck (region *R*_1_), the CR-MSD in general scales with a smaller exponent than the ballistic ordinary MSD, but is still super-diffusive. This again indicates perturbation of local cellular order there. The configuration of smallest diameters, with a constant CR-MSD, represents an exception. As the bottleneck’s diameter in this case corresponds to only one or two cell diameters, cells are in a locked-in configuration as soon as they enter the narrow part so that movement relative to neighbours is bounded. In the context of thermal colloids, a similar effect is referred to as “single-file diffusion”. [64]

In summary, we observe that the narrow funnel forces cells to adopt a different arrangement. This results in different dynamics, not only inside the bottleneck, but also in the region ahead of it.

### 3.5 Varying the diameter gradient in the funnel region

Because of the large influence of the funnel, i.e. the transition region between wider and narrower parts, has on the collective order and dynamics of the cells, we also considered channels where this transition is smoother; in other words, the funnel is longer. In Fig. 6A, the velocity in channel direction *v_x_*(**r**, *t*) is compared for both cases. Velocities are higher for the more slender geometry in the region *R*_0_ in front of the bottleneck, but very similar in the other areas. This indicates that backflow is considerably reduced (see also Supp. Figs. 10,11) and that transport along the channel is enhanced. These findings are in line with the results for the scaling of the MSDs (Fig. 6B, Supp. Figs. 13,14): While in general the results are similar, in region *R*_0_ the ordinary MSD’s scaling exponent is closer to two for the slender geometry. In contrast, the exponents for the CR-MSD are smaller in regions *R*_0_ and *R*_1_, corresponding to reduced mixing of cells and more stable configurations. Not much difference can be observed in the positions of the maxima in the angular distribution of the RDF *f_ϕ_*(*ϕ*) (Fig. 6C, Supp. Fig. 12), while the typical nearest-neighbour distance *l*_max_ is lower for the slender geometry in regions *R*_1_ and *R*_2_, i.e. within and beyond the bottleneck. These findings for the MSDs and the spatial organisation of cells are in line with the observation of a reduced backflow ahead of the channel’s narrow section, which allows cells to squeeze through the bottleneck more efficiently. All in all, the more slender channel entry allows cells to stack more compactly and stably in the narrow region, and thus facilitates higher transport through the channel, which also leads to less backflow and congestion.

**Figure 6:**
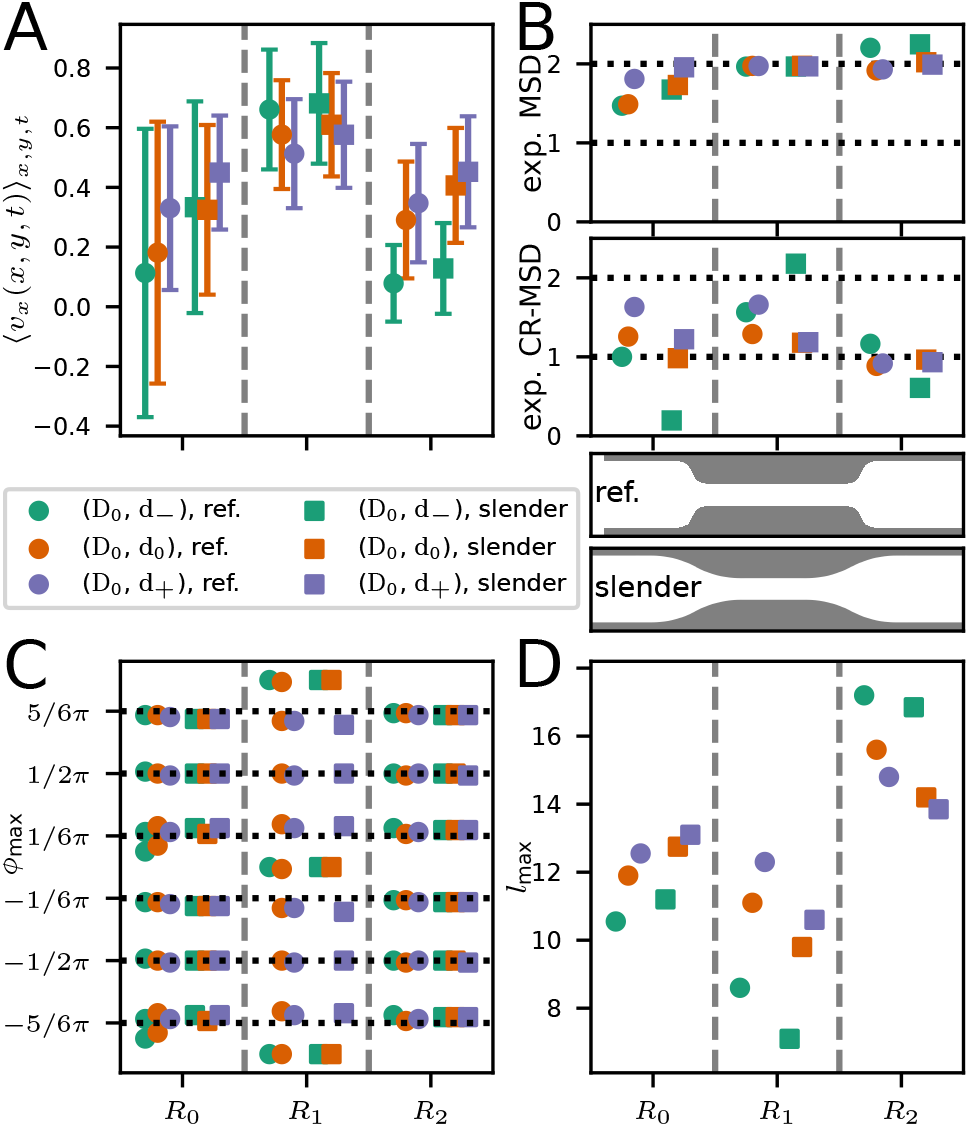
Comparison of internal organisation and dynamics for different designs of the channel entrance and exit. The reference design has sharper transitions while these are less abrupt for the “slender” geometry, as the sketches between panels B and D illustrate. (A) Average velocity in channel-axis direction in the three different regions. The higher values in region *R*_0_ reflect smoother transitions due to the reduced backflow. (B) Comparison of scaling of MSDs. The ordinary MSD (upper panel) scales very similarly for both, the CR-MSD’s exponent is generally lower for the slender geometry, which points to a more regular dynamics reduced relative motion of cells. (C) Angles at which maxima in the RDF appear. Results are very similar: the difference for (*D*_0_, *d*_0_) is attributable to the non-linear character of the maxima detection rather than a real difference in the RDF. (D) Typical distance of maxima in the RDF from the centre. For the same values for (*D, d*), the slender geometry leads to larger distances, i.e. lower cell densities, in region *R*_0_ and higher densities, or smaller distances, in the other regions. This is compatible with the observation that the slender geometry yields less congestion at the entrance and facilitates a denser packing inside the bottleneck.

In addition to variations in channel geometry, the system’s behaviour can be changed by variation of parameters. In general, the influence of parameters on the dynamics of single cells and collectives in the CPM is examined in detail in Ref. [22]. In the context of our study, cell-cell adhesion has an especially interesting effect. Higher cell-cell adhesion energies cause cells to follow their neighbours more eagerly at the edge of the epithelial sheet. This leads to an increase in mean cell velocities and densities and thus facilitates faster invasion (see section A.4).

## 4 Conclusions

In this work, we have used computer simulations of a Cellular Potts Model (CPM) which emulates individual cells that exhibit internal polarisation and pairwise, as well as cell-substrate, interactions. We aimed at investigating the influence of a bottleneck channel on an advancing cellular monolayer. To this end, we compared the local spatial organisation of the cellular agglomerate and the collective as well as individual dynamics of cells ahead of, within and beyond the bottleneck across varying geometries. The radial distribution function, the mean velocity of cells in the direction of the channel axis, and the scaling of the cells’ mean-squared displacement served us as the corresponding observables.

While cells in an unconstricted channel tend to organise in sixfold-coordinated hexagonal structures and move along the channel with only minor variations in speed and density, the presence of a bottleneck leads to accumulation and reflux of cells near the entrance to the narrow part of the channel, and to distortions of hexagonal order. This effect strongly depends on the relative diameters of the broader and tighter channel sections as well as the design of the transition region in between: The larger the dimensions of the geometry, the less effect on order and dynamics in the cell sheet, while more “streamlined” transitions from broad to narrow facilitate a tighter packing of cells in the bottleneck and hence reduce backflow and pile-up at the bottleneck entrance. These effects, especially on the spatial organisation of cells in the interior of the epithelial sheet, are particularly strong when bottleneck dimensions approach the diameter of only a few cells.

From the physics perspective, the cells show interesting collective properties. On the one hand, we observe features reminiscent of a compressible fluid: increase in density and flow-velocity in the narrow part of the channel while the CR-MSD shows that single particles are not caged completely. On the other hand, there is short-range hexagonal order, which is a typical attribute of solids. The cell sheet thus displays features of both solids and fluids, as it represents a collective composed of ballistically moving agents. In accordance with previous studies, it thus behaves like a glass [59, 60, 61, 62, 58, 63] that oozes through down the channel. These collective properties are an emergent phenomenon of the interaction of our model cells via cell-cell adhesion which leads to strong coherence in the monolayer and dictate its properties.

The focus of our work lies on the internal structure and single-cell dynamics inside the sheet and how it is influenced by a bottleneck. Previous works, both experimental [9, 10, 11, 12, 17, 13, 14, 16] and numerical [25, 42], have concentrated on velocity and density fields or the shape of the invasion front in channels of constant [9, 10, 11, 12, 17, 13, 14, 16, 25, 42] or linearly changing [17, 25] diameter. In many of these studies strong density fluctuations in the channel are reported, a feature that our model did not reproduce. Additional intercellular coupling mechanisms, possibly of the polarisation field, may be required to reproduce this phenomenon.

From the biologist’s point of view, under the assumption that our simplistic model captures the system’s essential properties, our observations provide information on how an epithelial cell-sheet might react when confronted with physically constricting environments, for example during cancer metastasis or morphogenesis [2, 65]. It would be interesting to see if our results can be reproduced in laboratory experiments. As our analyses heavily rely on positional data of cell centres, potential experiments to test our predictions need to be able to identify and track individual cells or nuclei. Under a more general perspective, the work presented here provides an additional test case on the question of whether the CPM, despite its relatively simple structure, is specific enough to capture variations in the behaviour of cells under varying experimental conditions and can therefore serve as a useful tool for the understanding of cellular dynamics.

In addition, the versatile nature of our model makes it readily adaptable to reflect more closely the conditions of certain experimental systems that for example use a particular type of cells, substrate or channel geometry. This can be achieved not only by variation of parameters, but also by further modifications, such as the use of a viscoelastic substrate [44], additional interaction mechanisms, or different confining geometries. Also more detailed models for the internal dynamics could be introduced. For example, replacing the scalar polarisation field with a polar or nematic field could provide a means of mimicking the symmetry properties of the stress fibres in the cytoskeleton and the resulting anisotropic tensions.

Note also that there is no “genetic” component to the model, as all cells behave identically. This in particular excludes the possibility of investigating the interplay of distinct cell types, for example the consequences of differential adhesion [66, 34] where cell-cell interactions differ according to cell type. It also does not allow for the emulation of leader cells, which are cells that more aggressively invade empty spaces and drag other cells with them. This effect has been investigated thoroughly in experiments as well as numerical studies [17, 67, 25, 68, 69] and leader cells are assumed to play an important role in cell migration. As such, the implementation of different cell types would increase its versatility even more and allow for the investigation of additional biologically relevant phenomena.

## Acknowledgements

E.F. and F.K. would like to thank Sophia Schaffer, Matthias Zorn, and Joachim Rädler for stimulating discussions. This work was funded by the German Excellence Initiative via the program “Nanosystems Initiative Munich” (NIM) and by the Deutsche Forschungsgemeinschaft (DFG, German Research Foundation) - Project-ID 201269156 - Collaborative Research Center (SFB) 1032 - Project B2.

The preprint template has been taken from https://github.com/brenhinkeller/preprint-template.tex/

## Supplementary Materials

### A Model and parameters

We use the model as presented in [22] with a slightly adapted cell cycle and parameters. The source code and all necessary input data are available upon reasonable request.

The constriction can in principle and in conjunction with active cell motility lead to a large accumulation of cells, which is equivalent to a compression of individual cells. To ensure a sufficiently fine discretisation of each cell even under such a compression, we increased the individual cell area roughly by a factor of 4 compared to our previous confluent tissue simulations in [22]. To that end, we increased the average polarization field *∊*_0_ and at the same time decreased the area stiffness parameter *κ_A_*.

Furthermore, we increased the effective temperature to allow for larger cell size fluctuations. We kept the signalling radius at the same absolute value, which implies that the ratio between signalling radius and cell diameter decreased. We slightly increased the cell-cell adhesion energy. Finally, we decreased cell motility compared to [22] by decreasing cell polarisability and increasing the membrane stiffness parameter.

#### A.1 Simulation procedure

Before the actual simulation, cells are randomly seeded in a reservoir left of the channel entrance. For a certain burn-in time (700 time steps in our case), cells can evolve freely in this reservoir. In this time, the cell density grows to maximum density in the reservoir. After that time, cells are finally allowed to enter the channel and the actual simulation starts. Also then, cell division is limited to the reservoir.

#### A.2 Implementation of cell cycle

Compared to [22], we modified the quiescent phase in that we made the transition to growth stochastic in order to achieve a less deterministic cell-cylce and to avoid unwanted synchronisation of cell division. The rest remains unchanged.

A freshly initialised cell, or a daughter cell right after cell division, is in the *quiescent state*. In the original model, the cell would now switch into the *growing state* if its size, via fluctuations, grew larger than *rA*_ref_ (for explanation of parameters see Table 1 in the next section). In the version used for this work in contrast, the cell switches into an intermediate state, the *refractory state*, in which it does not grow. From there, it can switch into the growing state in every simulation step with a probability of *p* = 1 − exp(−1/*T_q_*) with *T_q_* a newly introduced characteristic waiting time. From there, the cell cycle works exactly as described in [22].

**Table 1:** Parameter values used in our study, following the the notation convention from [22].

| Cell parameters |  |  |  |
| --- | --- | --- | --- |
| Parameter | Value | Function | Comment |
| $\epsilon_0$ | 75 | strength of cell-substrate adhesion<br>/average polarisation speed | |
| $\Delta\epsilon$ | 45 | cell polarisability: the cell polarisation $\epsilon$ is in the interval $\epsilon \in [\epsilon_0 - \Delta\epsilon/2, \epsilon_0 + \Delta\epsilon/2]$ | parameter influences the persistence of cells and overall invasion speed |
| $D$ | 0 | strength of cell-substrate dissipation: energy penalty for abandoning a tile, i.e. breaking cell-substrate bonds | finite values slow down dynamics |
| $\mu$ | 0.10 | cytoskeletal update rate | sets overall timescale. |
| $R$ | 2 | signalling radius | when a cell conquers a tile, the polarisation field increases for tiles of this cell within radius $R$ of the new tile |
| $\kappa_A$ | 0.10 | area stiffness | |
| $\kappa_P$ | 0.20 | perimeter stiffness | together with $\kappa_A$ and $\epsilon_0$ sets the preferred size of a single cell $A_{\text{ref}}$:<br>$A_{\text{ref}} = (\epsilon_0 - 2\pi\sqrt{3}\kappa_P) / (\sqrt{3}\kappa_A)$ |
| $B$ | 10 | strength of cell-cell adhesion | See A.4 for results for $B = 5$. |
| $\Delta B$ | 0 | strength of cell-cell dissipation | |
| Parameters for cell division |  |  |  |
| $T_g$ | 350 | cell growth duration | in this time, the cell grows exponentially to double its size by increasing the preferred cell-size by adjusting $\kappa_A$ and $\kappa_P$. |
| $T_d$ | 50 | cell division duration | in this time, the cell is immotile and the polarisation relaxes to neutral. After that, the cell splits |
| $T_q$ (new) | 100 | cell quiescence duration | after cell division, the cell does not grow for this time on average |
| $r$ | 0.50 | relative threshold for cell growth | after the quiescent phase, the cell starts growing if its size is larger than $rA_{\text{ref}}$ |
| Numerical parameters |  |  |  |
| $N_0$ | $\approx 1100$ | initial number of cells | ... |
|  | 700 | burn-in time | ... |
| | $D = D_0/2 : 3000,$<br>$D = D_0 : 4000,$<br>$D = 2D_0 : 5000.$ | simulation time | ... |
| | $2500 \times 1200$ | dimensions of simulation area | geometries available on request |
| $k_B T$ | 5 | effective temperature | reference energy of fluctuations in the Monte-Carlo algorithm |
|  | 1 | seed | seed for random number generator |

#### A.3 Parameters used

As we have used the model from [22], we use the notation convention for parameters from there. The values are given in Table 1. Corresponding configuration files and files containing geometry information are available upon request along with the code.

#### A.4 Variation of cell-cell adhesion

In this section we examine the effect of variations in the cell-cell-adhesion *B*. The cell-cell-adhesion defines the effective free energy gained from an edge that a cell shares with another cell compared to the situation where the edge borders to a free tile. A higher cell-cell-adhesion parameter thus favours cohesion of cells.

Interestingly, higher cell-cell adhesion also leads to faster dynamics in the invasion of the channel, as can be seen in Supp. Figs. 1A and 2A. There we show simulations with cell-cell adhesion parameter *B* = 5 in comparison to the simulations with *B* = 10 in the main text. Lower cell velocities for *B* = 5 are particularly prominent in regions *R*_1_ and *R*_2_ for medium and large bottleneck diameters *d*_0_, *d*_−_. One explanation is that for higher benefits through adhesion, trailing cells are faster to follow leading cells as they are faster to form common edges. In that way, advancements of a single cell are translated more directly into advancements of the collective.

While the peak-distribution in the angular distribution *f_ϕ_*(*ϕ*) is not significantly altered (Supp. Figs. 1C, 2C), the radial distribution *f_r_*(*r*) shows slightly smaller cell-cell distances for larger cell-cell adhesion (Supp. Figs. 1D, 2D) which additionally promotes cell-flux and thus faster invasion (Supp. Fig. 2A). The single-cell dynamics in the cell sheet scale comparably (Supp. Figs. 1B, 2E).

**Supplementary Figure 1:**
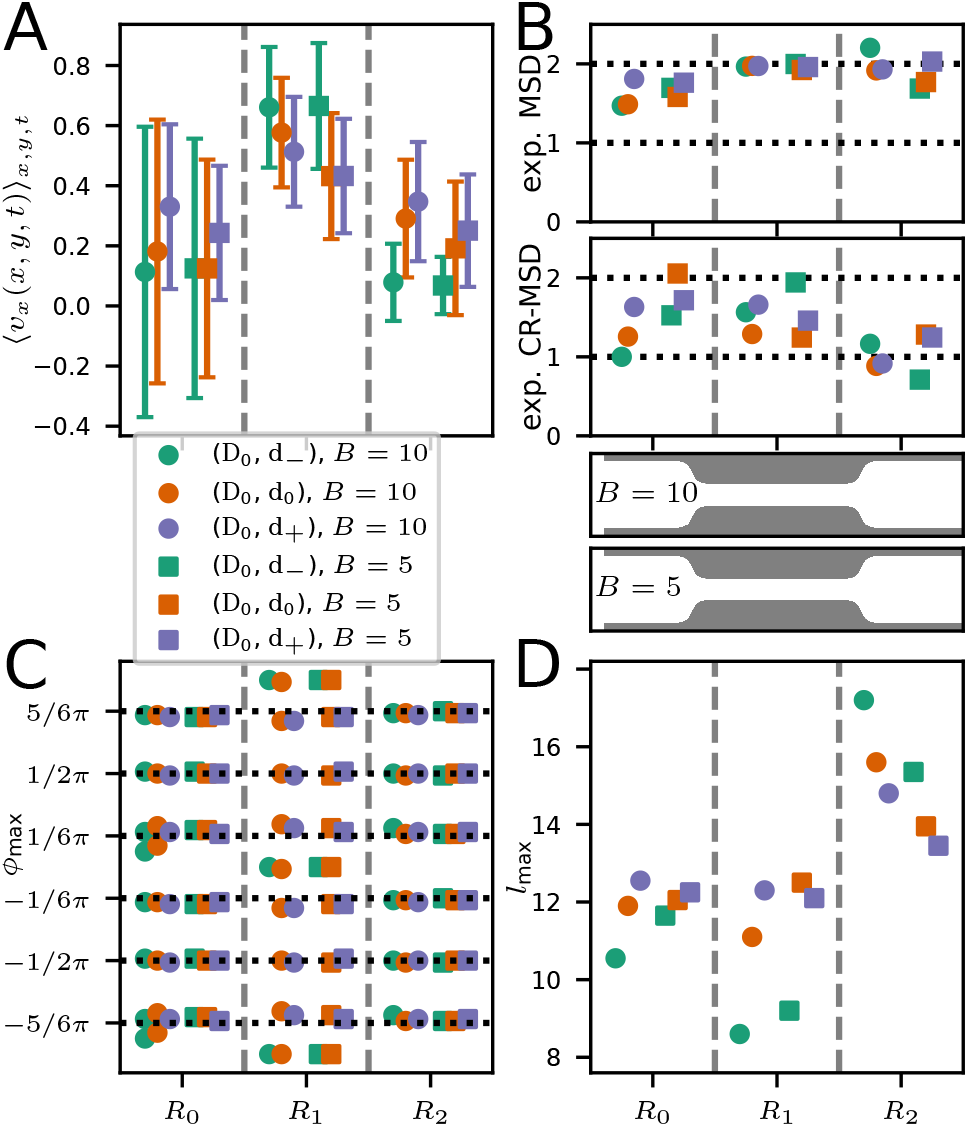
Comparison of structure and dynamics for different values of cell-cell adhesion energies. (A) Average velocity in channel-axis direction in the three different regions. Higher cell-cell adhesion leads to higher values, especially in region *R*_1_. (B) Comparison of scaling of MSDs. The ordinary MSD (upper panel) scales very similarly for both, showing ballistic motion in the narrow region and behind as well as reduced scaling as a sign of backflow before the bottleneck. (C) Angles at which maxima in the RDF appear. Results are very similar. (D) Typical distance of maxima from the center in the RDF. As can also be seen in Supplementary Figure 2, the cell-cell distance is larger for smaller cell-cell adhesion, especially in smaller channels. Due to the slower overall invasion, there are significantly less cells in region *R*_2_ for *B* = 5 than for *B* = 10. Hence cell-cell distances are higher there

**Supplementary Figure 2:**
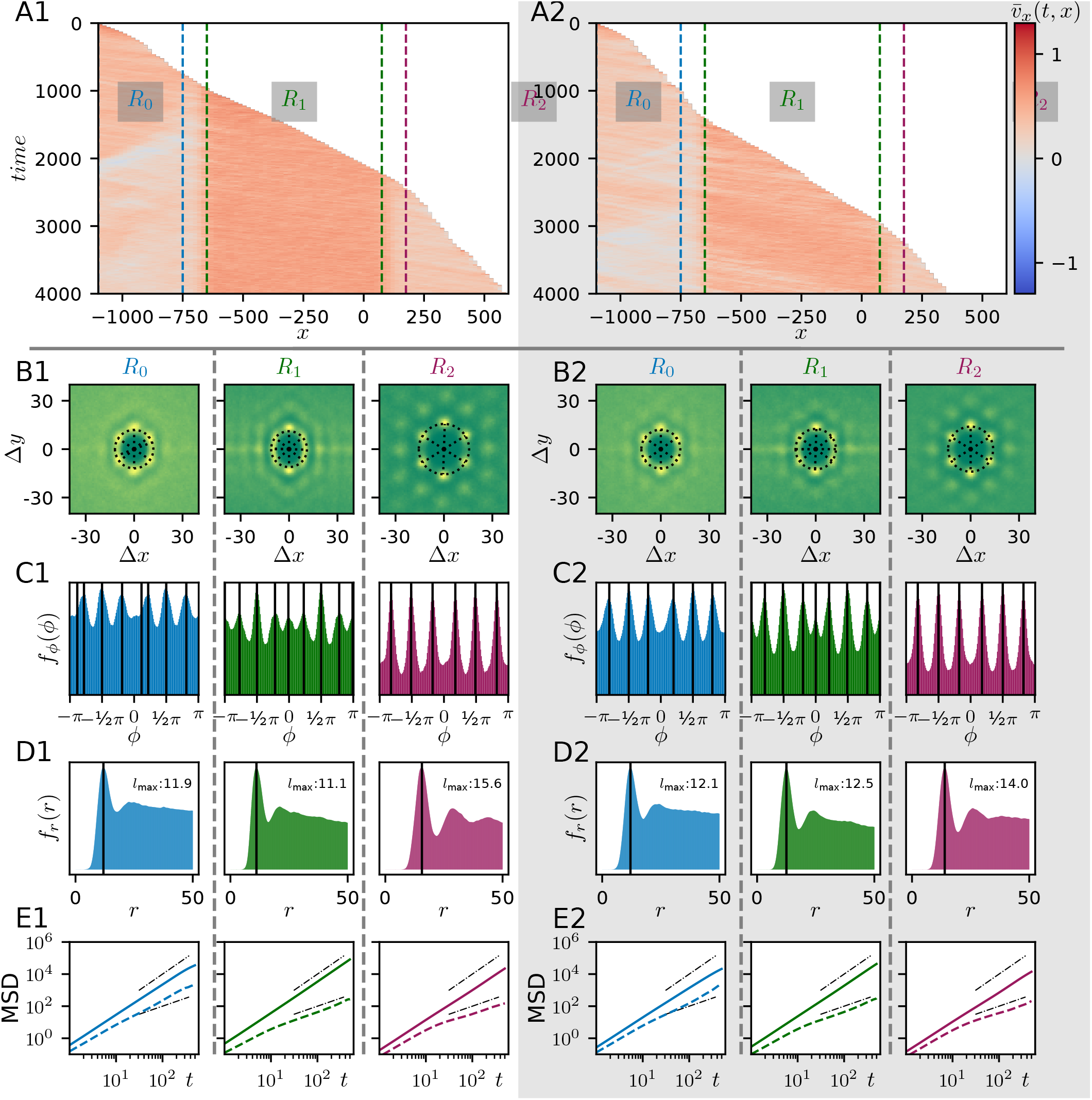
Comparison of order and dynamics for different values of cell-cell adhesion energy. Left side (panels Xa): B = 10, right side (panels Xb): B = 5). (A) Kymograph of the velocity in channel direction *v_x_*(*t, x*). (B-D) Radial distribution functions (RDFs) for the different regions of the channel. The angular position of peaks is mostly unaltered (panel C), while the distance between cells in larger for smaller adhesion (panel D). (E) Mean-squared displacement (solid line) of cells as well as cage-relative mean-squared displacement (dashed line). Black dashed-dotted lines show linear and quadratic scaling, respectively.

### B Methods

#### B.1 Analysis of the radial distribution function (RDF)

As explained in the main text, the RDF is defined as (Eq. (1) in the main text)

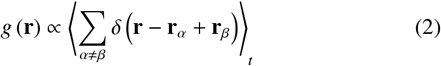

with the double sum running over all pairs of cells in the area under observation, **r**_*α*_, **r**_*β*_ being the positions of the cells [51], and the angular brackets indicating a time average.

The arrangement of peaks on a circle around the origin (see Fig. 4 in the main text) suggests to analyse the angular distributions of peaks on this circle as well as its radius. To this end, we define the radial and the angular distribution functions as:

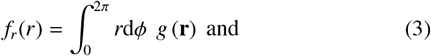

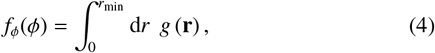

respectively. Here, polar coordinates are defined as **r** = (*r* cos(*ϕ*), *r* sin(*ϕ*)). Due to the symmetry *g* (**r**) = *g* (−**r**), *f_ϕ_*(*ϕ*) = *f_ϕ_*(*ϕ* ± *π*). The upper boundary for the radial integration is *r*_min_, the position of the first minimum after the first maximum of *f_r_*(*r*); this choice ensures that only the nearest neighbours contribute to *f_ϕ_*(*ϕ*). From *f_r_*(*r*), we obtain *l*_max_ simply as the maximum closest to the center, and *r*_min_ the minimum following it.

For the analysis of *f_ϕ_*(*ϕ*), we aim at detecting the number of peaks or, more specifically, judging how well it fits a regular six-fold symmetry or whether the peaks at 0, ±*π* are more prominent.

##### B.1.1 Maxima detection

For the peaks, we on purpose chose a simple definition. A peak is defined as a maximum *ϕ*_max_ if *f_ϕ_*(*ϕ*_max_) > 0.5 *f_ϕ_*(*ϕ*_max,0_) with *ϕ*_max,0_ the absolute maximum of *f* (*ϕ*). Also, no maxima are allowed closer than 0.1*π* to another maximum to avoid small bumps on the flanks of peaks in *f_ϕ_*(*ϕ*) to be counted. Still, sometimes additional peaks are detected, as in Fig. 3 Bb in the main text.

Also, as we are looking at an observable with a hard criterion (maximum/ no maximum), gradual changes in the distribution lead to jumps in the results. This is why in the region where the distribution is transitioning form six-fold to two-fold, it should not be overestimated when maxima are detected at *ϕ* = 0, ±*π*. To show that the method is nevertheless suitable to detect qualitative changes in the system, we will have a detailed view on the angular distributions in the next section.

##### B.1.2 Detailed view on angular distributions

In Supplementary Figure 3, we can look at the distributions for parameters *D* = *D*_0_, *d* ∈ {*d*_−_, *d*_0_, *d*_+_}, the examples that are also shown in the centre column of Fig. 4. We also show fits of a distribution with two and six peaks, respectively. For that, we use superpositions of the so-called wrapped Cauchy-distribution [70]:

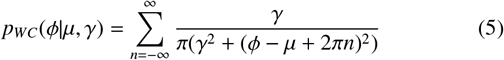

where *γ* is the scale factor and *μ* is the peak position of the “unwrapped” distribution. *γ* is the only fitting parameter, positions of the peaks are set to ±*π/*6, ±*π/*2, ±5*π/*6 and 0, ±*π*, respectively.

Supplementary Figure 3 shows that the algorithm finds maxima at ±*π/*6, ±*π/*2, ±5*π/*6 as well as at 0, ±*π* only for the *D*_0_, *d*_−_ configuration in region *R*_1_. In the other cases, the algorithm agrees with what is obvious to naked eye.

In addition, the similarity of the distributions can be estimated using the L-2 norm

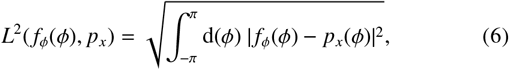

where in our case we compare the measured distribution *p*(*ϕ*) ≡ *f_ϕ_*(*ϕ*) to a fitted distribution of six regularly distributed peaks *p*_6_(*ϕ*), a fitted distribution of two peaks *p*_2_(*ϕ*), and a flat distribution *p_flat_*(*ϕ*). Results are shown and discussed in the lower panel of Supp. Figure 3. From what we observe there, we can conclude that the approximation with a distribution with two or six peaks captures the structure of the distributions sufficiently well and that in particular, one fits the data better than the other in all cases considered.

#### B.2 Mean-squared displacement (MSD) and cage-relative mean-squared displacement (CR-MSD)

The ordinary mean squared-displacement (MSD) is defined as follows:

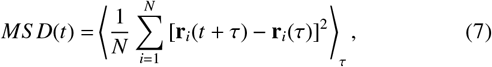

with *N* the number of cells under considerations and **r**_*i*_(*t*) the position of cell *i*. In analogy the cage-relative MSD (CR-MSD) is defined as [57, 58]

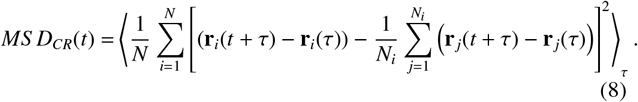

Here, the second sum runs over all neighbours of cell *i* at time point *τ*, *N_i_* is then umber of neighbours at of cell *i* at time *τ*. Hence, the CR-MSD compares displacement of cells to the displacement of their original neighbours. The neighbours are determined by Voronoi tesselation.

We calculate the MSD for the last 500 time steps of the simulation and determine the scaling exponent *x* of the MSDs by computing

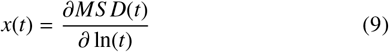

and averaging over *x*(*t*) for the last half of these 500 time steps. This approach assumes a structure of the MSD as *MS D*(*t*) ∝ *t*^*x*^.

The CR-MSD stresses the local, or relative, dynamics. In a solid, the CR-MSD was to saturate reflecting “caging” of the particles, while the ordinary MSD also picks up global movement of the system as we have due to the invasion dynamics. A CR-MSD scaling linearly means that particles move diffusively relative to each other.

**Supplementary Figure 3:**
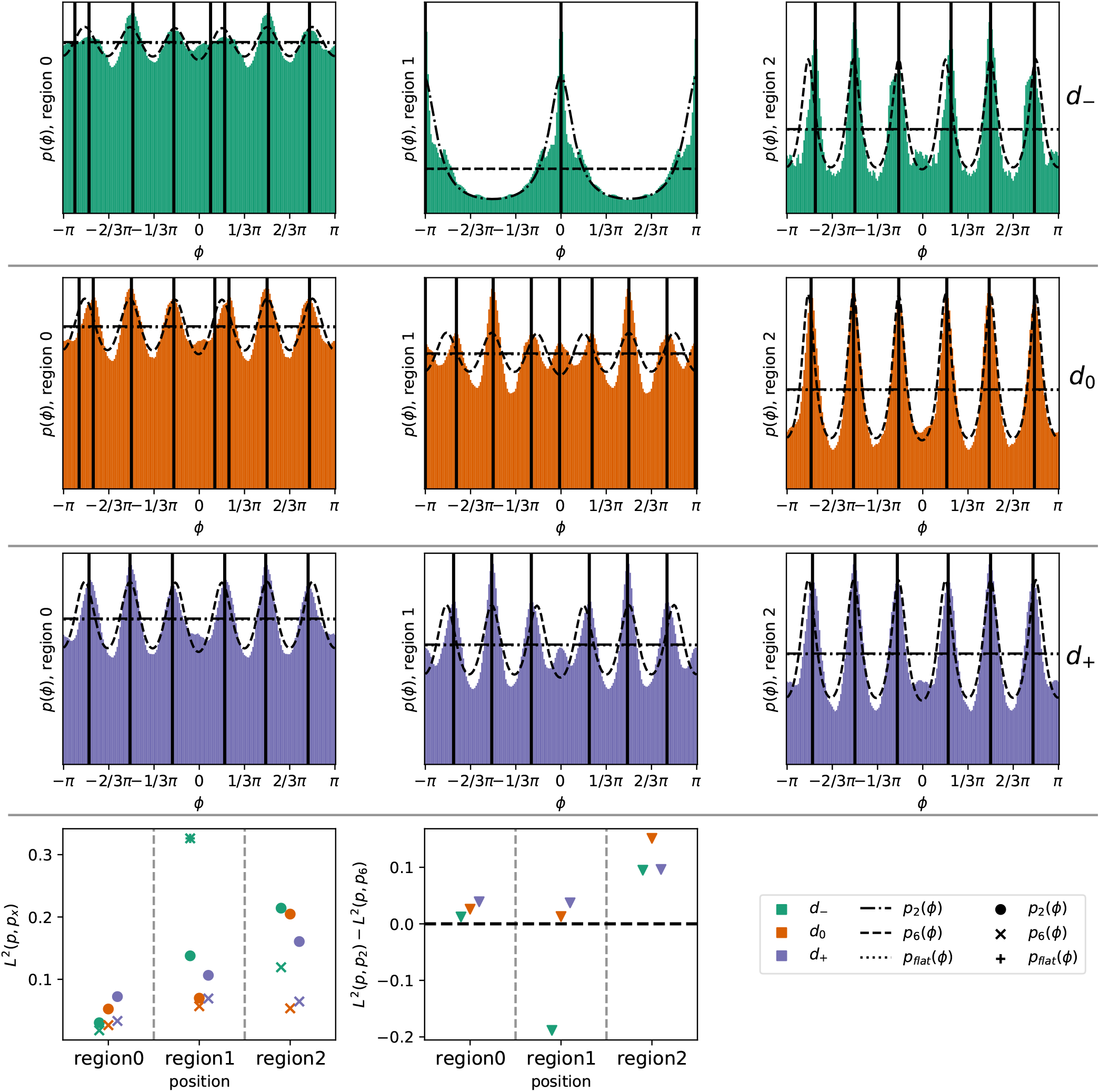
Comparison of measured marginalised radial distributions f_ϕ_(ϕ) to theoretical distributions. Upper three rows show data for parameter sets (*D*_0_, *d*_−_), (*D*_0_, *d*_0_), (*D*_0_, *d*_+_) for the three regions separately. Data is in colours, approximations as different lines according to legend. While in region *R*_2_, for all parameter combinations, we see six regularly distributed peaks. In region *R*_0_ for all three case as well as in region *R*_1_ for parameters *d*_0_ and *d*_+_ peaks are less prominent, but still sixfold and regular. Only for region *R*_1_ and the narrowest bottleneck diameter *d*_−_, there are only two peaks at 0, ±*π*. This is also reflected in the lowest row, where the *L*^2^-distances between *p*(*ϕ*) = *f_ϕ_*(*ϕ*) and *p_z_, p*_6_, *p_flat_* are shown. It becomes clear from the right plot that *p*_6_ is the better fit except for (*D*_0_, *d*_−_) in region *R*_1_. . Actually, the best fit for the distribution less less close to the data, turns out to be identical to the flat distribution.

### C Supplementary figures

#### C.1 MSDs and CR-MSDs under variation of D and d

Supp. Fig. 4 shows complete MSDs and CR-MSDs. The corresponding scaling exponents are plotted in Figure 5 in the maintext.

**Supplementary Figure 4:**
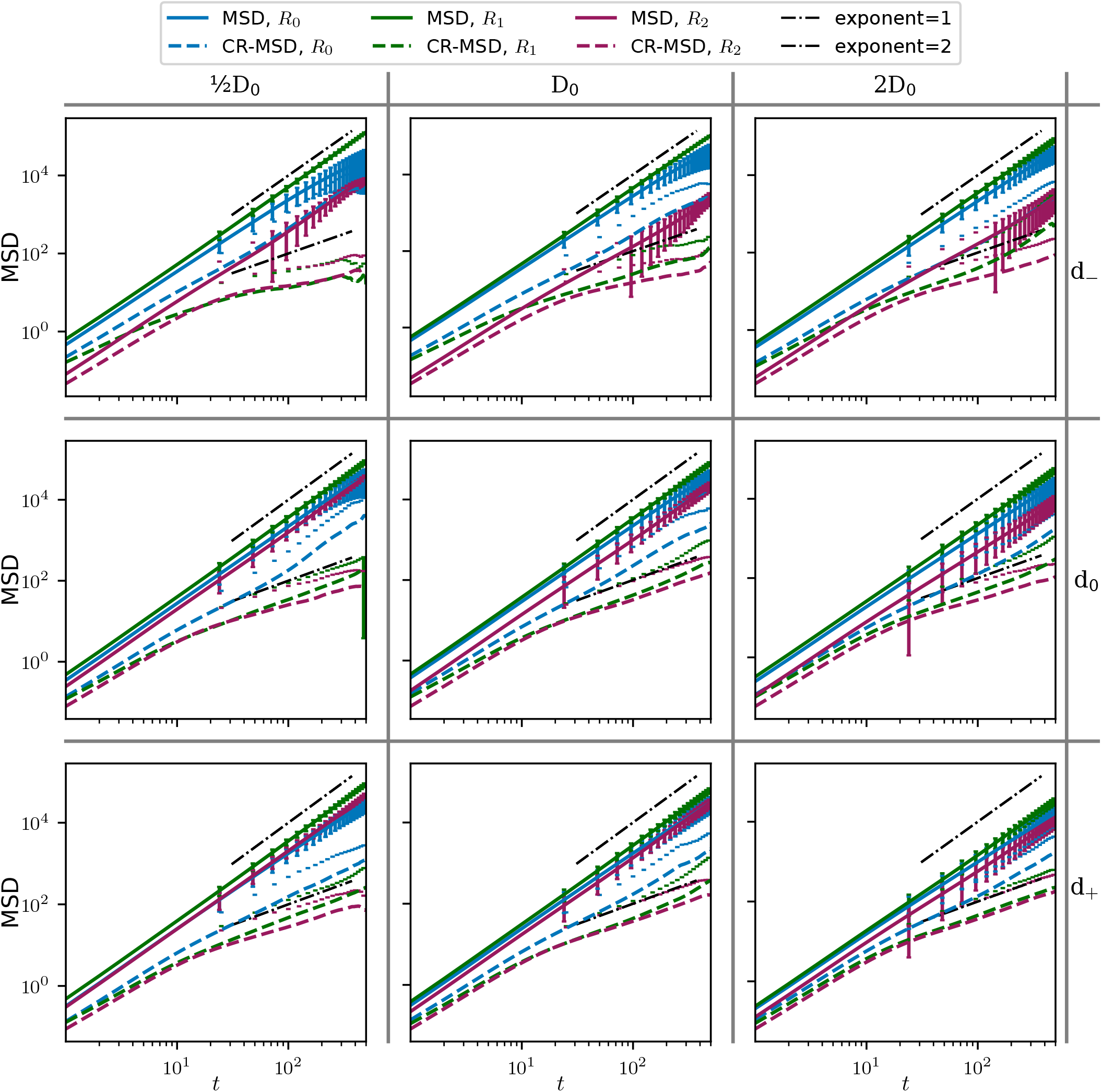
Log-log plot of MSD and CR-MSD for varying D, d. The solid lines show the ordinary MSD, the dashed lines the CR-MSD. The ordinary MSD scales with exponent 2 in all regions, while the CR-MSD after an initial period of ballistic scaling grows to slower from *t* ≈ 10 on. The CR-MSD in region *R*_0_, shown in the blue dashed lines, is an exception to this, for all parameter configurations it grows faster than the other CR-MSDs, which is caused by the backflow in this region.

#### C.2 Additional figures for the configuration with a straight channel

Supp. Fig. 5 shows snapshots of density and velocity fields for an arbitrary time point. It is the corresponding figure to Fig. 2 for a bottleneck channel in the main text. In Supp. Fig. 6, we compare the structure of the cell sheet for varying diameters *D*. It corresponds to Fig. 4 for bottleneck channels in the main text.

Results for MSDs and CR-MSDs in straight channels are shown in Supp. Fig. 7 (analogous to Supp. Fig. 4) and Supp. Fig. 8 which is the same as Figure 5 in the maintext, but for straight channels of different diameters.

A summary of structural and dynamical differences is given in Supp. Fig. 9.

**Supplementary Figure 5:**
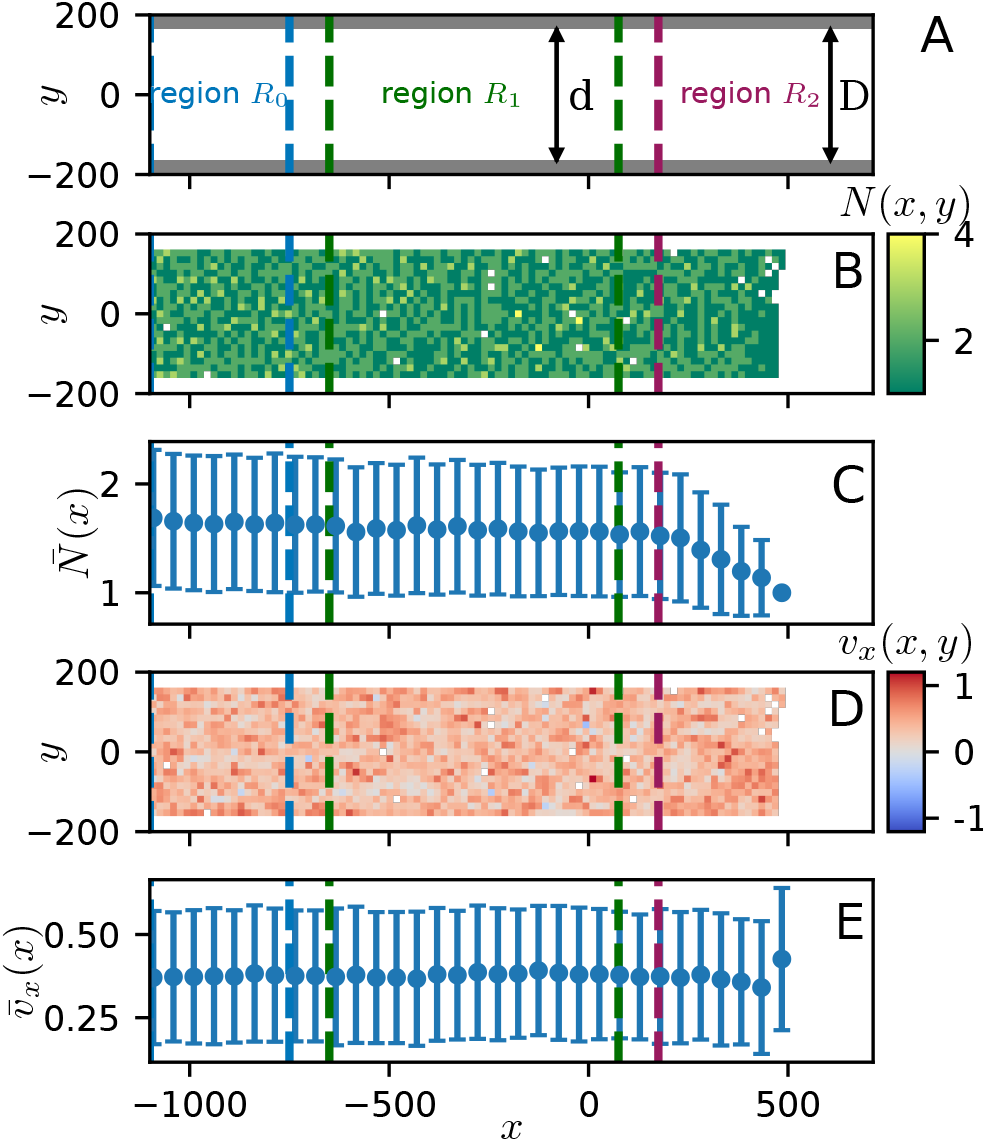
Example for simulation in a channel without bottleneck. Widht D = D_0_ according to the convention in the main text. Plot analogous to Fig. 2 in the main text. (A) Geometry, (B) heatmap for particle number, (C) mean number of particles per bin. Apart from a slightly less dense region at the invasion front, the density is constant. (D) heatmap for velocity in channel direction, (E) mean velocity in channel direction, the value is constant throughout the cell-sheet.

**Supplementary Figure 6:**
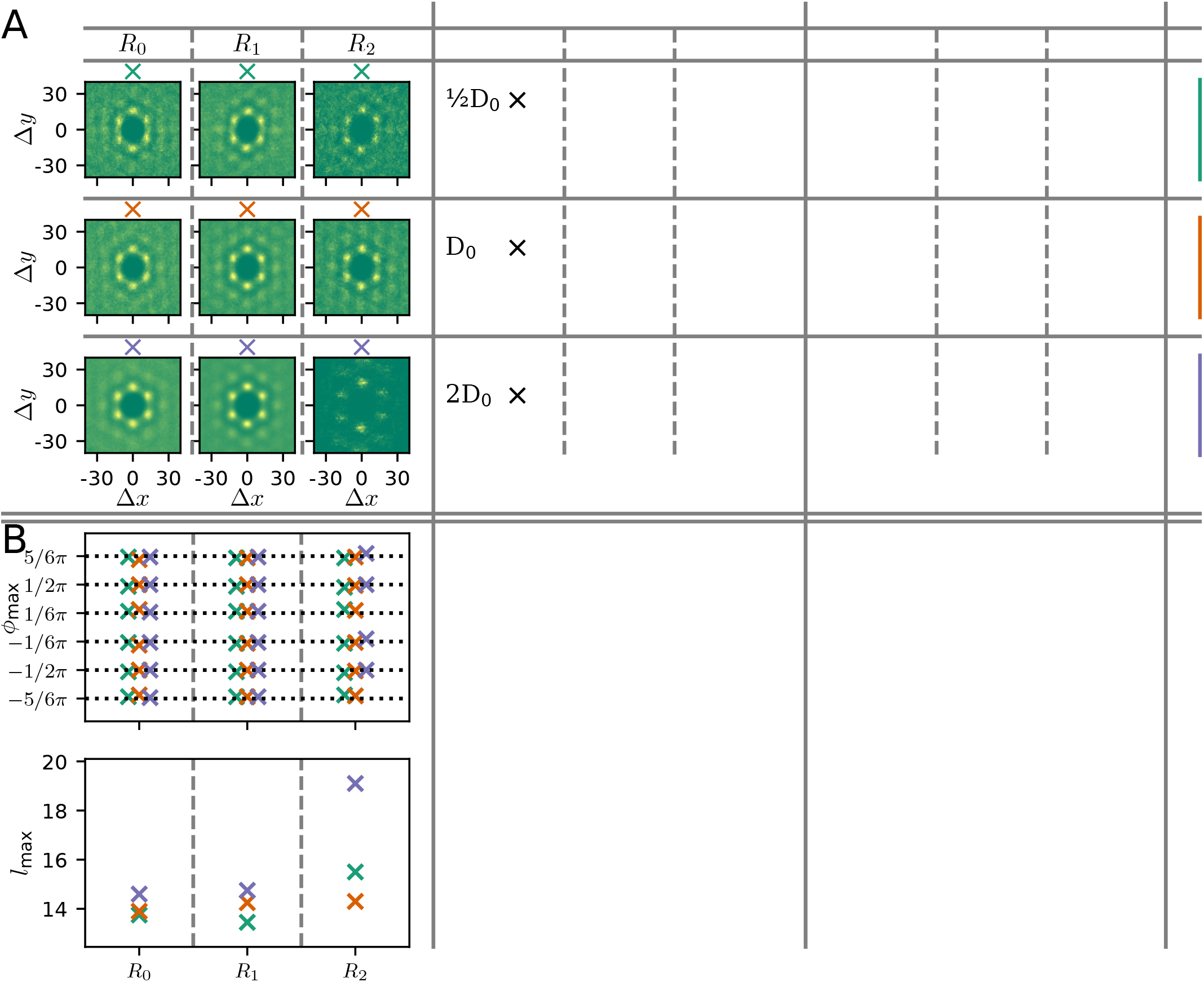
Comparison of cellular order in straight channels of varying width. (A) Radial distribution functions (RDF) for different diameter *D* for the different regions along the channel. From top to bottom: increase in *D*. In all regions, cells arrange in a regular hexagonal configuration. Low contrast in region *R*_2_ for *D* = 2*D*_0_ is due to the reduced invasion speed in the broader channel. (B) Results of quantitative evaluation of the results in part (A). The region is marked by the position of the markers along the horizontal axis of the respective plots. Varying colors indicate variations in *D* as the small symbols above the plots in part A symbolise. *Upper panel*: Positions of maxima in the angular distribution obtained from radial integration of the RDF. *Lower panel*: position of maximum in the radial distribution obtained from the angular integral of the RDF. Distance of cells is a bit higher closer to the front, but very similar in regions *R*_0_ and *R*_1_.

**Supplementary Figure 7:**
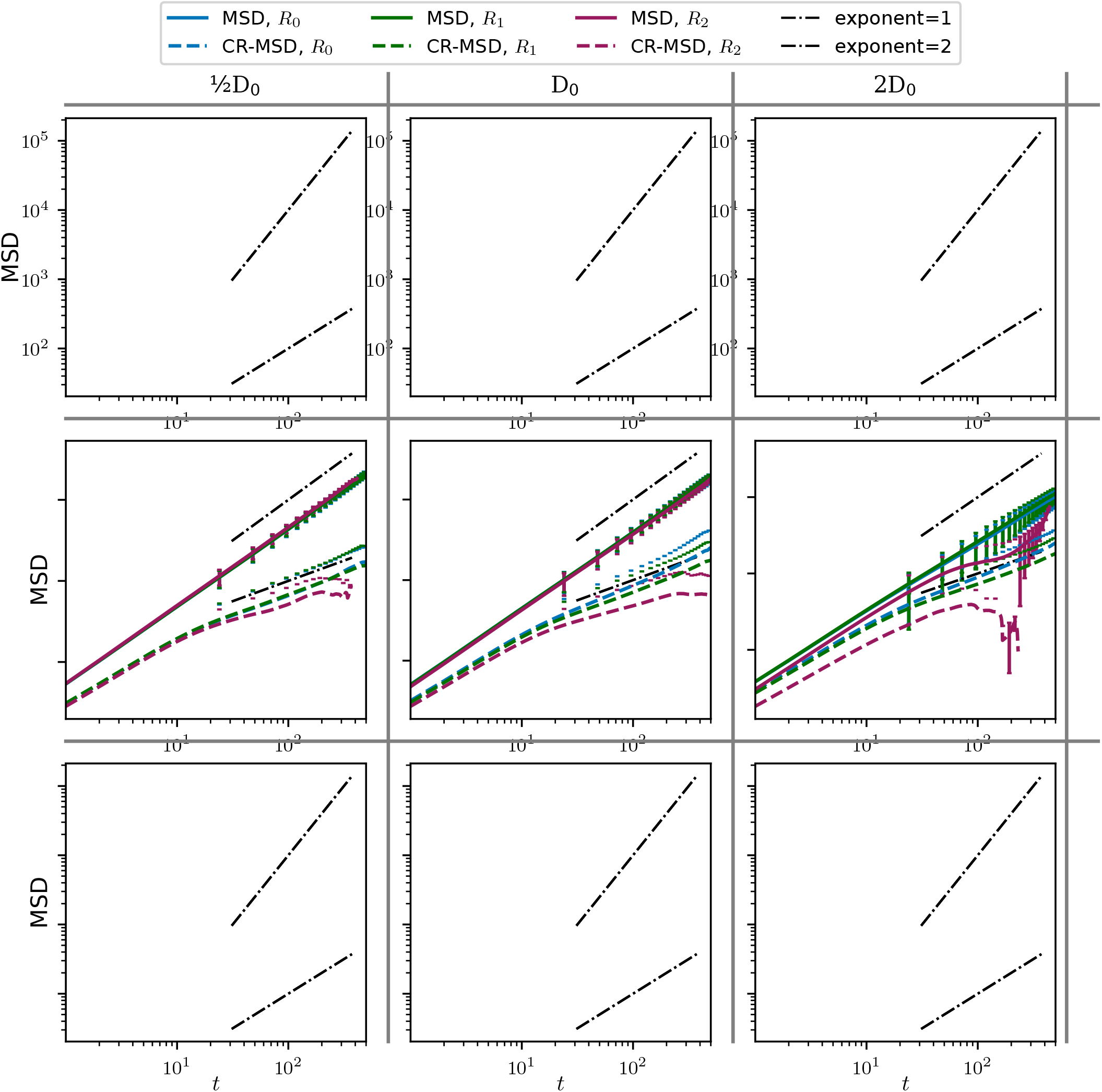
Log-log plot of MSD and CR-MSD for varying D in a straight channel (d = D). The solid lines show the ordinary MSD, the dashed lines the CR-MSD. Ordinary as well as cage-relative MSD behave very similar across regions *R*_2_0 and *R*_1_. In region *R*_2_, close to the front, the CR-MSD scales significantly lower, especially for *D* = *D*_0_. Compared to Supp. Fig. 4, there are no signs of backflow in region *R*_0_, as CR-MSD scales linearly.

**Supplementary Figure 8:**
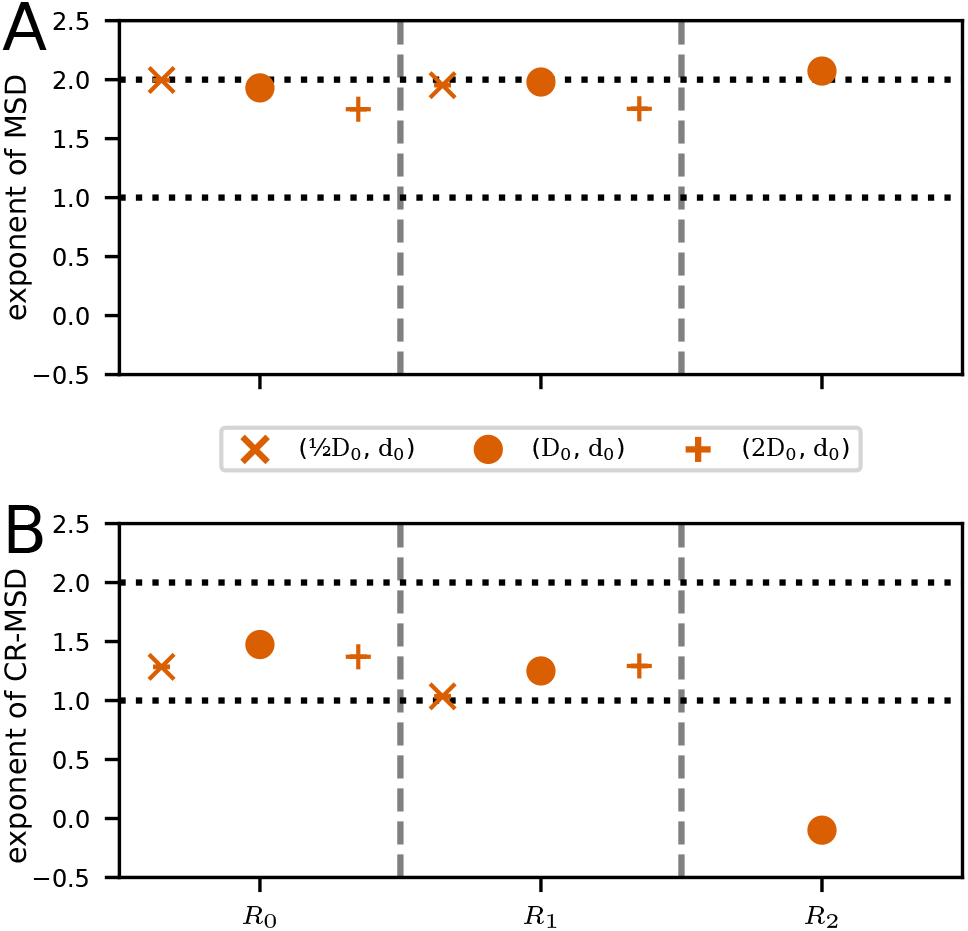
Comparison of MSDs in straight channels. (A) Exponent from monomial fit to the late MSD for separate for regions *R*_0_, *R*_1_, and *R*_2_. Results reflect what could already be seen in Supp. Fig. 7: the ordinary MSD in (A) expresses ballistically moving cells while the reduced exponents by the CR-MSD in (B) are lower. The stronger reduction in region *R*_2_ is due to the presence of the front.

**Supplementary Figure 9:**
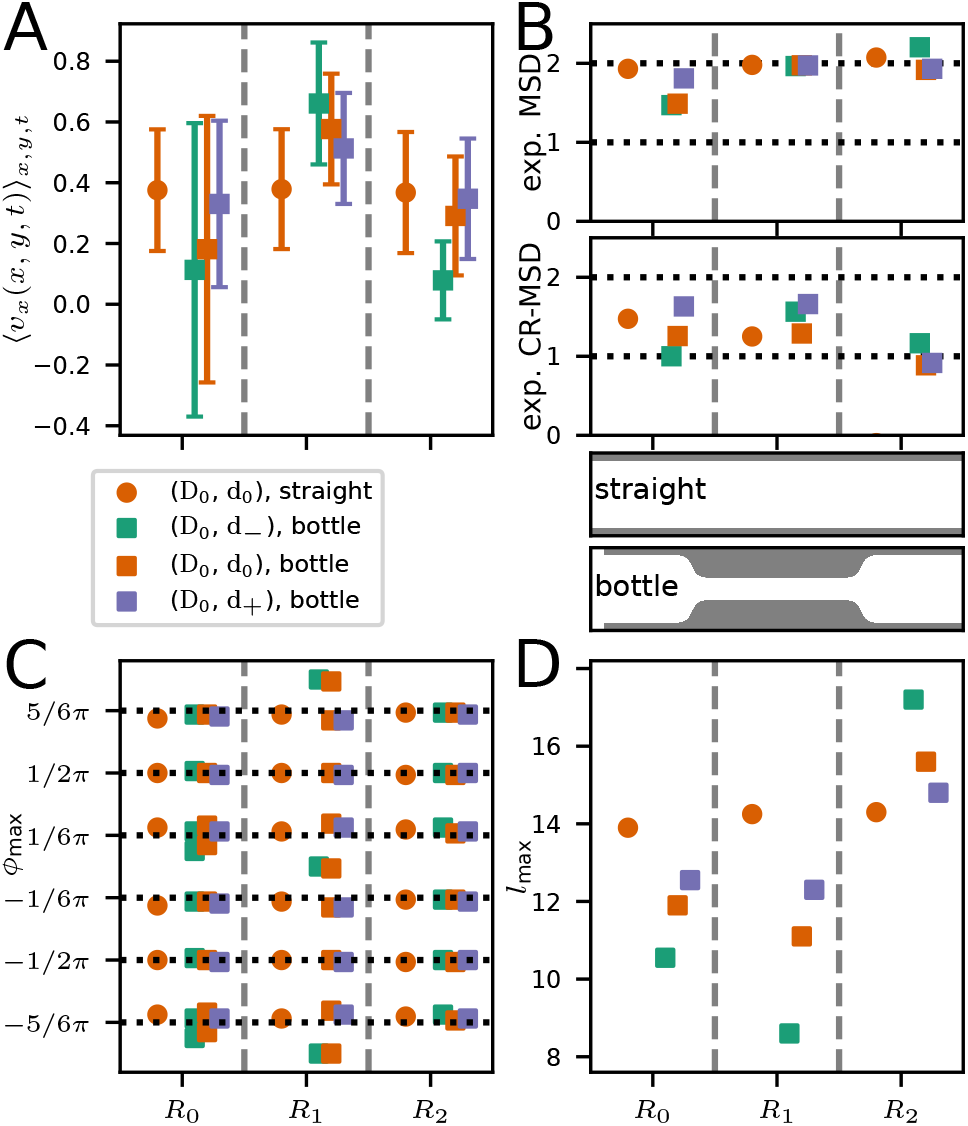
Comparison of internal organisation and dynamics for bottleneck channels and a straight channel as sketches between panels B and D illustrate. (A) Average velocity in channel-axis direction in the three different regions. (B) Comparison of scaling of MSDs. (C) Angles at which maxima in the RDF appear. (D) Typical distance of maxima in the RDF from the center.

#### C.3 Additional figures for the configuration with longer funnel/ smoother transition

For more detailed comparisons of results for different funnel shapes, we present additional plots. Supp. Figs. 10–14 are in essence the same figures as in the main text and Supp. Fig. 4, but for the geometry with the slender funnel.

**Supplementary Figure 10:**
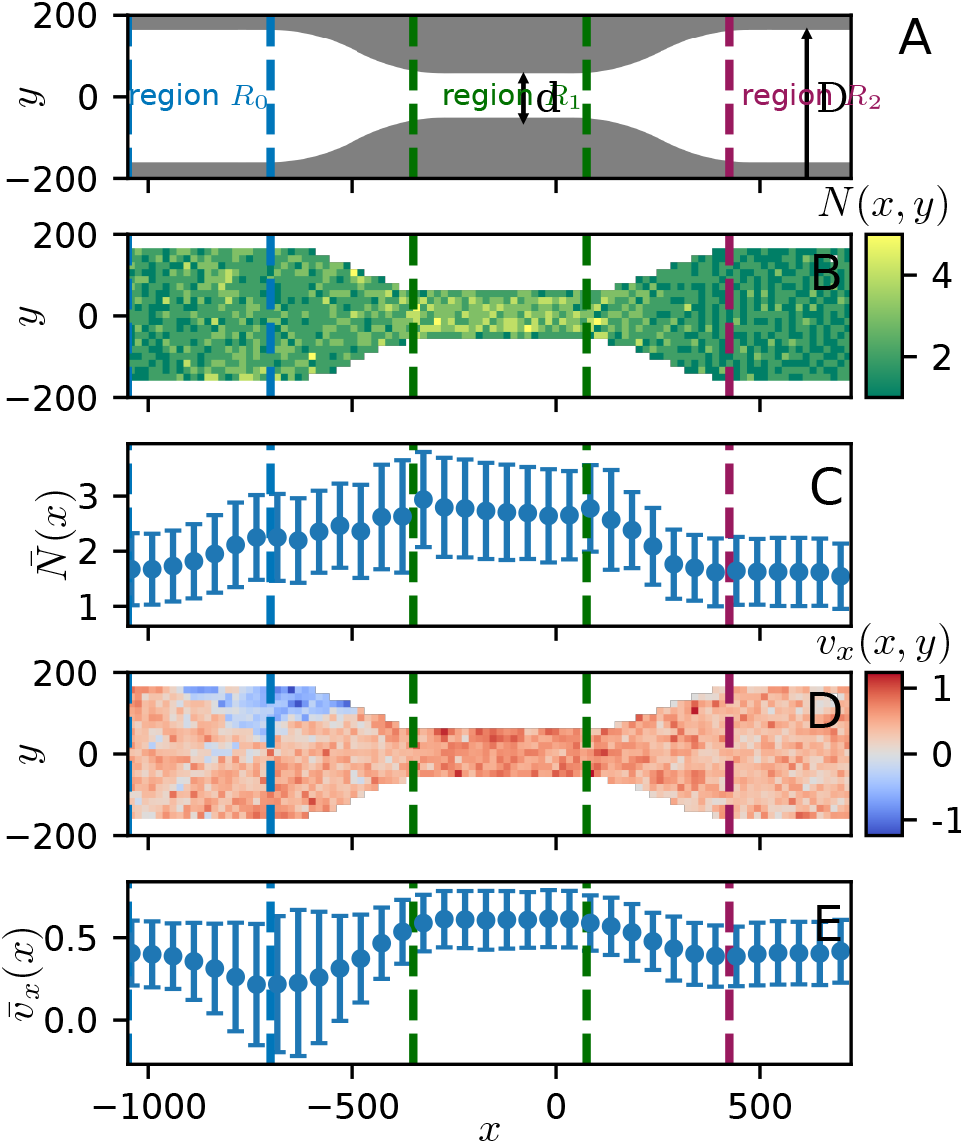
Example for simulation in a channel with longer funnel. Diameters D = D_0_, d = d_0_ according to the convention in the main text. Plot analogous to Fig. 2 in the main text. (A) Geometry, (B) heatmap for particle number, (C) mean number of particles per bin, (D) heatmap for velocity in channel direction, (E) mean velocity in channel direction. In comparison to Fig. 2, density and velocity are even more increased in the narrow part. The transition in density as well as velocity is distributed over the full lenght of the funnel, but backflow is still visible.

**Supplementary Figure 11:**
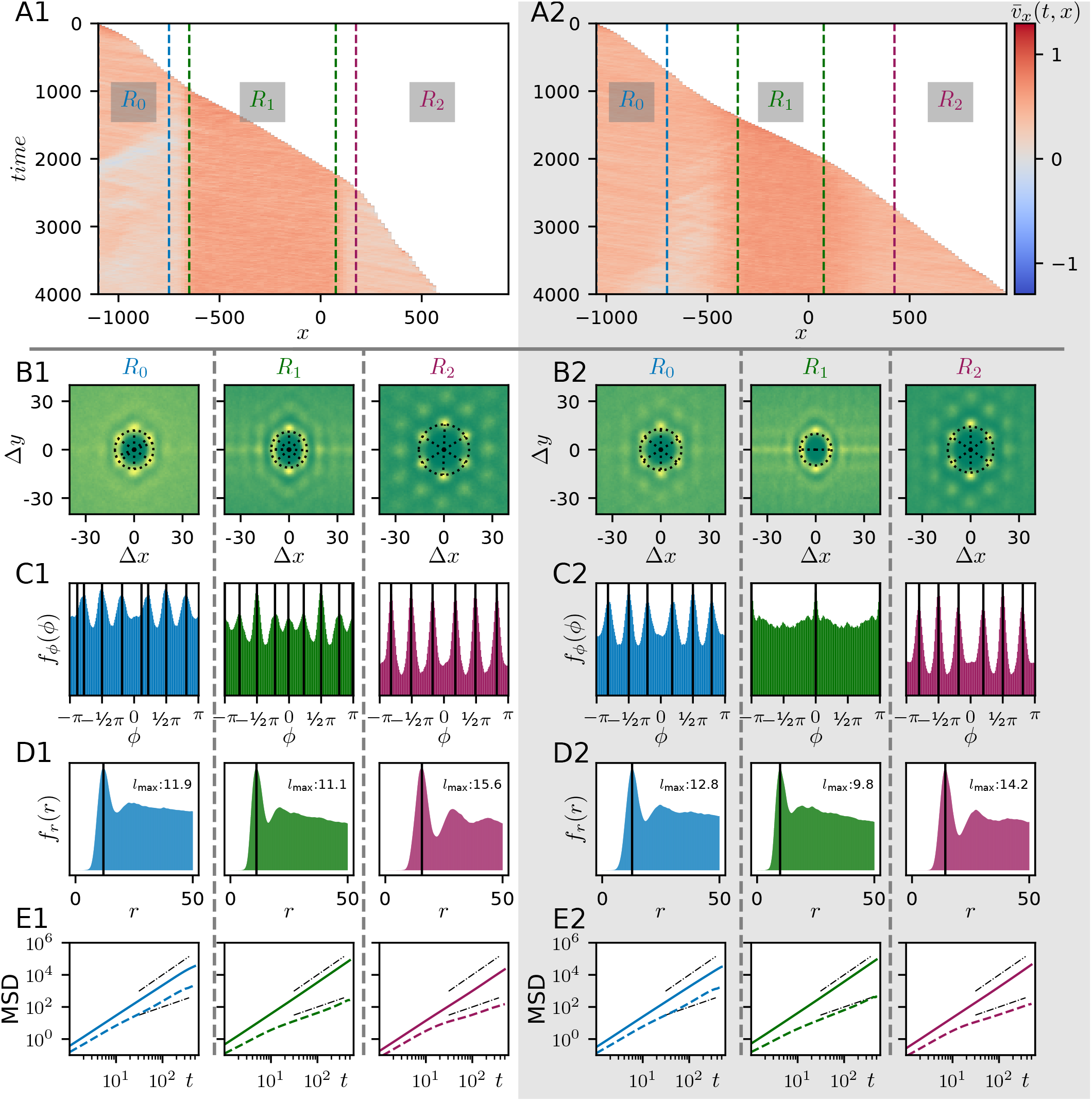
Comparison of order and dynamics for a constricted channel, parameters D = D_0_, d = d_0_. Left side (panels Xa): bottleneck with steep transition, right side (panels Xb): bottleneck with smoother transition. (A) Kymograph of the velocity in channel direction *v_x_*(*t, x*). Dashed lines indicate transition regions from wider to narrower part. In comparison, the slender funnel leads to later and less backflow and in conjunction an overall faster invasion as the steeper frontline, especially in region *R*_2_, shows. (B-D) Radial distribution functions (RDF) for the different regions of the channel. Panels C show the marginal distribution from integrating over the radius, and panels D integration over the polar angle. Vertical black lines in C mark maxima in the angular distribution as detected by the algorithm (for details see section B). The vertical black lines in D mark the respective maximum of the radial distribution which gives the typical nearest-neighbour distance. Especially in the narrow region *R*_1_, the slender transition creates a more dense cell sheet and different internal order as the angular distribution shows. (E) Mean-squared displacement (solid line) of cells as well as cage-relative mean-squared displacement (dashed line). Black dashed-dotted lines show linear and quadratic scaling, respectively.

**Supplementary Figure 12:**
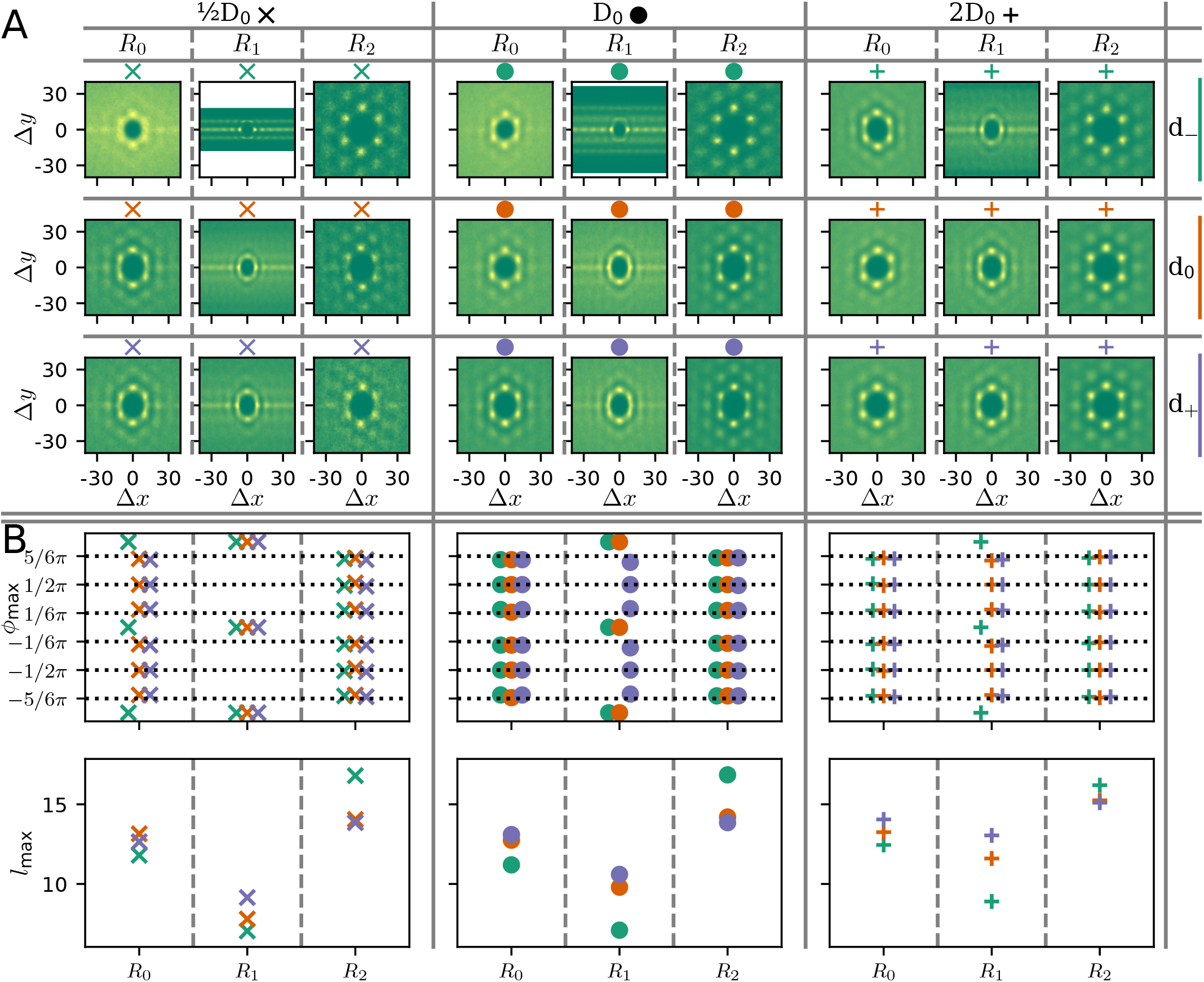
Comparison of cellular order in bottleneck channels of varying width with smoother transitions. (A) Radial distribution functions (RDF) for different diameter *D* and *d* for the different regions along the channel. From left to right: increase in *D*, from top to bottom: increase in *d*. (B) Results of quantitative evaluation of the results in part (A). The region is marked by the position of the markers along the horizontal axis of the respective plots. *Upper panel*: Positions of maxima in the angular distribution obtained from radial integration of the RDF. *Lower panel*: Position of maximum in the radial distribution obtained from the angular integral of the RDF. In comparison to the analogous data in Figure 4 in the main text, for the steeper transition, the order in region *R*_1_ is different for parameter sets *D* = *D*_0_, *d* = *d*_0_ and *D* = 2*D*_0_, *d* = *d*_0_. Especially there is no parameter set where the two distinct peak-patterns overlap. Also, variations in cell-cell distance/ cell density are stronger.

**Supplementary Figure 13:**
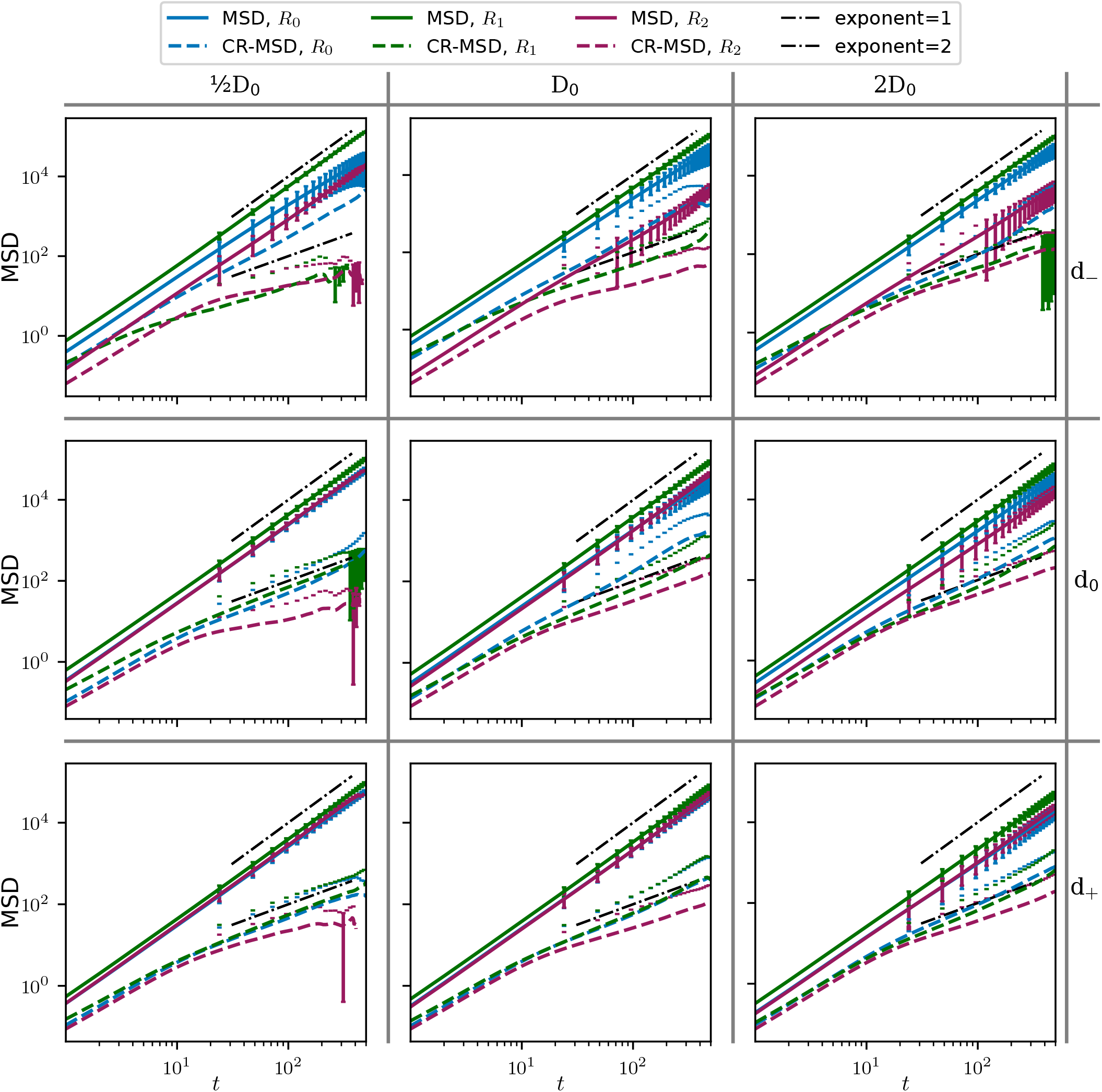
Log-log plot of MSD and CR-MSD for varying D, d in the channel with the smoother transition. The solid lines show the ordinary MSD, the dashed lines the CR-MSD. The ordinary MSD scales with exponent 2 in all regions, while the CR-MSD after an initial period of ballistic scaling grows to slower from *t* ≈ 10 on. The exception is again the CR-MSD in *R*_0_ that in channel regions of smaller *D* or *d* grows comparable to the ordinary MSD.

**Supplementary Figure 14:**
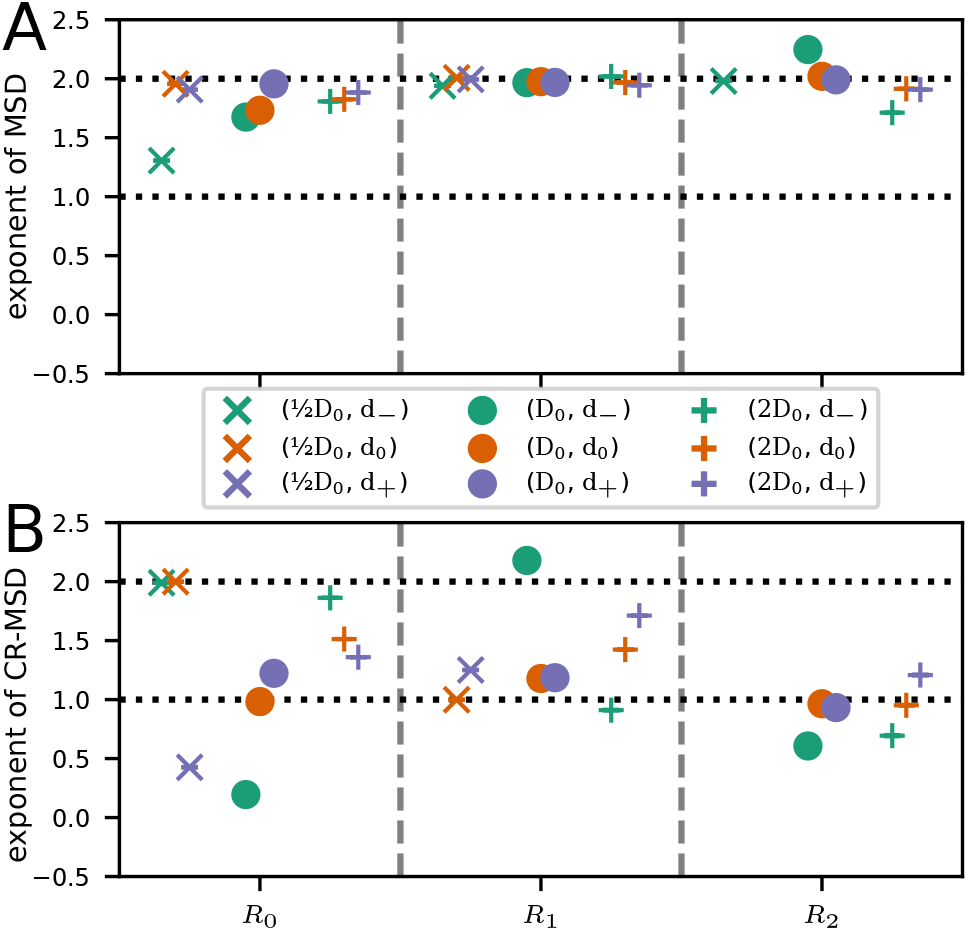
Comparison of MSDs in constricted channels of varying width with slender transition. (A) Exponent from monomial fit to the late MSD for same parameter configurations as in Supp. Fig. 13 and separate for regions *R*_0_, *R*_1_, and *R*_2_. (B) Corresponding exponents from monomial fits to the late CR-MSD. Except green cross and the green dot for parameters, the CR-MSD is consistently lower than the ordinary MSD. Missing data in region *R*_2_ is because invasion had not progressed significantly far into the region for a sufficiently long time.

